# Cell cycle exit is required for differentiation and cell fate diversification but not neoblast specialisation in planarians

**DOI:** 10.64898/2026.09.16.750956

**Authors:** Sophie Peron, Olena-Maria Balta, Addison Noronha, Toby Smillie, Vincent Mason, Alberto Pérez-Posada, Jordi Solana

## Abstract

In planarians, adult stem cells constantly proliferate and differentiate into dozens of different cell types to maintain tissue homeostasis. However, how cell fate acquisition is coordinated with cell cycle progression remains poorly understood. To address this, we generated single-cell RNA-sequencing datasets from cells enriched for 2C or 4C DNA content and from animals subjected to RNAi against the cell cycle regulators *cdh1* and *h2b*. We found that progenitor populations from multiple lineages contain actively cycling cells enriched in the 4C state. Cell fate acquisition to most cell types is acquired progressively during neoblast differentiation. Comparative analysis of cell cycle regulators confirmed a simplified regulatory repertoire in *Schmidtea*, supporting a central role for *cdh1* in controlling cell cycle exit. RNAi-mediated knockdown of *cdh1* caused homeostasis defects, impaired regeneration, accumulation of neoblasts and depletion of progenitors. Notably, specialised neoblast populations remained detectable despite continued cell-cycle progression, demonstrating that neoblast specialisation does not require cell-cycle exit. In contrast, progression from specialised neoblasts into progenitors and subsequent cell fate diversification was strongly impaired. Together, our findings distinguish two successive steps in planarian cell fate acquisition: neoblast specialisation can be initiated during active proliferation, whereas subsequent differentiation and fate diversification require *cdh1*-mediated cell-cycle exit.

## Introduction

Stem cells must be tightly regulated to ensure that the correct structures form at the appropriate location, size, and time during development, growth, and regeneration. Central to this regulation is the balance between self-renewal and differentiation. Although this balance is fundamental for tissue maintenance and regeneration, the molecular mechanisms that control it remain only partially characterised across animal species.

In planarians, growth, tissue homeostasis, and regeneration are sustained by a population of adult pluripotent stem cells known as neoblasts ^1^. These cells are the only proliferative cell type in the animal and are commonly identified by their high expression of *smedwi-1* ^2^. Neoblasts continuously proliferate and differentiate under homeostatic conditions to support normal tissue turnover, while retaining the capacity to generate every differentiated cell type ^3–5^. Following injury, neoblast proliferation increases dramatically through well-characterised mitotic peaks that drive regeneration ^6–8^. Likewise, feeding induces peaks of proliferation, whereas prolonged starvation is associated with a stable basal proliferation rate ^6,9^.

The neoblast population is transcriptionally heterogeneous, reflecting different stages of lineage commitment ^10,11^. Several studies have shown that subsets of neoblasts express distinct fate-specific transcription factors (FSTFs) during both homeostasis and regeneration ^12^. More recently, single-cell transcriptomic analyses have identified multiple transcriptionally distinct neoblast classes corresponding to different cell fates ^11,13–16^. These fates are only achieved after successfully exiting the cell cycle, as neoblasts lose *smedwi-1* expression and differentiate into specialised cell types. However, how neoblasts maintain the appropriate balance between proliferation and cell cycle exit/differentiation in these dynamic contexts is only beginning to be understood.

Neoblasts give rise to a wide variety of differentiated cell types. This process could occur through progressive fate restriction over multiple cell cycle or through fate acquisition within a single cell cycle followed by differentiation and maturation after cell cycle exit. There is recent evidence for a single step model of differentiation in which neoblasts progressively acquire lineage identity during the S/G2/M phases of a single cell cycle ^3^. Indeed, FSTF expression is less frequent during G1 compared to later cell cycle phases ^3^. Neoblasts expressing FSTFs can divide asymmetrically, producing one daughter cell that remains pluripotent and ready to express different FSTFs on the next cycle, and another that exits the cell cycle and differentiates ^3^. Neoblast specification is likely to be irreversible, since experimentally blocking the production of progenitors by supressing FSTFs results in smaller colonies ^17^.

Not all cell fates are specified directly within cycling neoblasts. Investigation of the transcription factors expressed in neoblasts subclusters and early progenitors proposed that lineage diversification occurs both within cycling neoblasts, after cell cycle exit within post-mitotic progenitors depending on the lineages ^14^. Despite these advances, it remains unclear how the decision to exit the cell cycle is coordinated with lineage commitment and differentiation, representing one of the major outstanding questions in planarian stem cell biology.

Planarians have lost many of the genes involved in vertebrate cell cycle regulation ^18^. Notably, they lack key components of the spindle assembly checkpoint, including MAD1 and MAD2, as well as several regulators of CDK activity required for S-phase entry. In *Dugesia japonica*, Sato and coworkers ^19^ identified *cdh1* as the only known regulator of CDK activity encoded in the genome. RNAi-mediated knockdown of *cdh1* resulted in neoblast hyperproliferation and impaired differentiation into epithelial, intestinal, and muscle lineages. These findings indicated that *cdh1* is required for cell cycle exit and established that cell cycle exit is essential for the successful differentiation of neoblasts ^19^.

Although previous studies have provided fundamental insights into neoblast biology, most have examined stem cell behaviour in non-physiological contexts, including enzymatic dissociation, sublethal irradiation, or transplantation. Particularly, fate specification in neoblasts was investigated in the context of regeneration by King et al ^14^. Here, we investigate the relationship between cell cycle progression and differentiation in *Schmidtea mediterranea* during homeostasis. To capture stem cell states as close as possible to their physiological condition, we generated single-cell transcriptomic datasets using ACME fixation and SPLiT-seq ^20,21^, incorporating fluorescence-activated cell sorting (FACS) to enrich for intact single cells with either 2C or 4C DNA content. We further combined these datasets with RNA interference of the cell cycle regulators *cdh1* ^19^ and *h2b* ^22^ to examine the consequences of disrupting cell cycle progression.

Our analyses show that progenitor populations across multiple lineages contain actively cycling cells and that lineage commitment occurs at different stages depending on the cell type in uninjured animals. Comparative analyses reveal that *Schmidtea* possesses a simplified repertoire of canonical cell cycle regulators, consistent with a central role for *cdh1* in controlling cell cycle exit. Our results distinguish two successive steps in planarian cell fate acquisition: neoblast specialisation, which occurs independently of cell-cycle exit, and subsequent progenitor differentiation and fate diversification, which require *cdh1*-mediated cell-cycle exit. Together, these findings provide new insights into how cell cycle regulation is coordinated with lineage commitment and differentiation in planarian stem cells.

## Results

### A multiplex integrated single-cell study to investigate planarian cell cycle and exit

To better understand the regulation of the planarian cell cycle and its relationship with the different neoblast classes and clusters, we generated two scRNA-seq datasets using ACME and SPLiT-seq ^20,21^. In the first experiment, we leveraged the FACS sorting step to isolate in one library cells with 4C DNA content to enrich G2 neoblasts (Figure 1A). Two further libraries contained 2C DNA cells, and an extra library contained 2C-4C cells together as regularly done. In the second SPLiT-seq experiment, we multiplexed ACME dissociated RNAi cell samples after knocking down *h2b*, as it is known to deplete neoblasts and early progenitors ^22^, and *cdh1*, the *Schmidtea mediterranea* ortholog of a gene described to inhibit cell cycle exit in the planarian species *Dugesia japonica* ^19^ (Figure 1B). We multiplexed two biological replicates of each RNAi, along with control RNAis in one plate, and obtained six sublibraries with 2C-4C cells.

**Figure 1:**
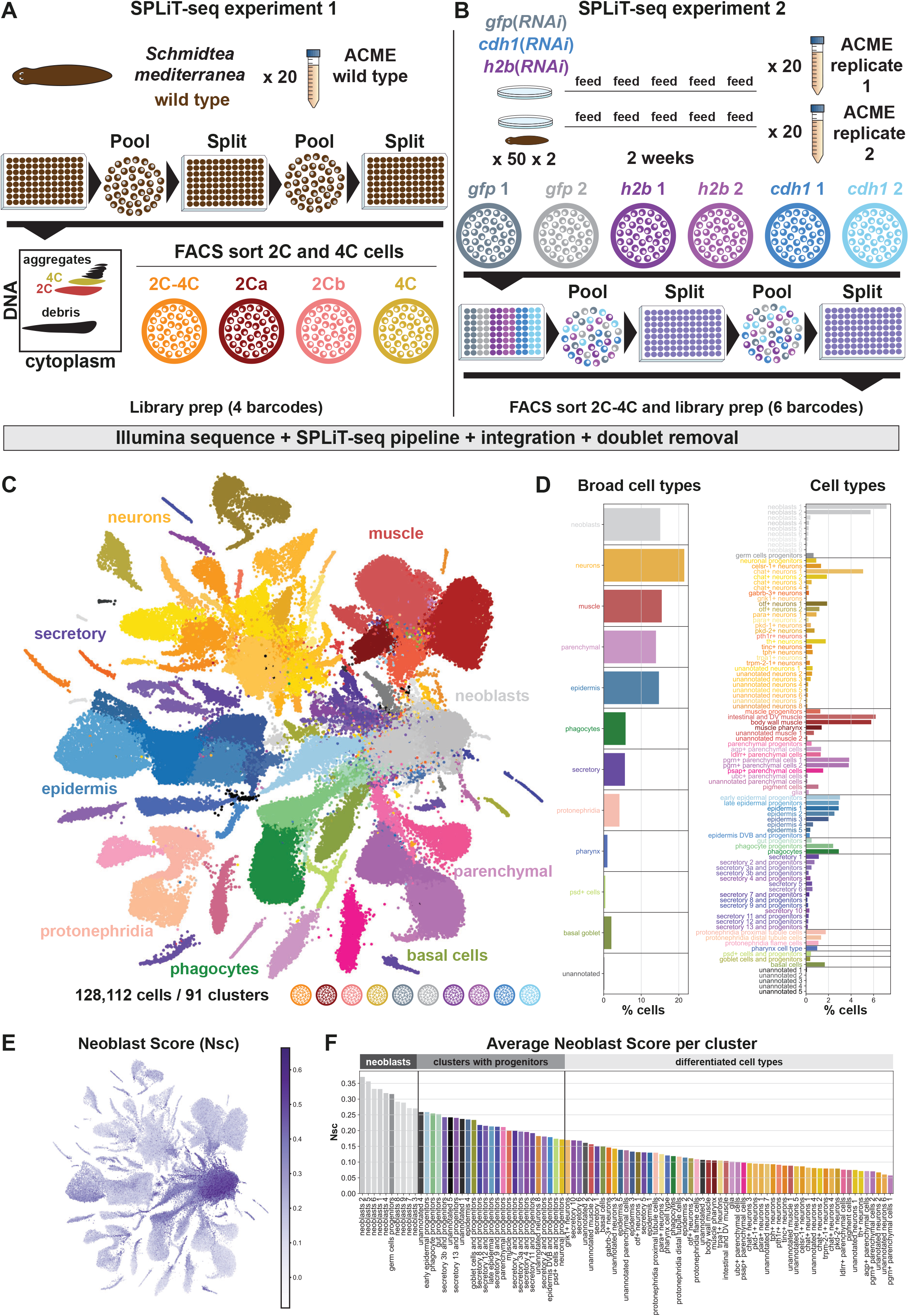
A multiplexed integrated single-cell dataset to investigate planarian cell cycle and exit. A. Experimental design of the SPLiT-Seq experiment 1. B. Experimental design of the SPLiT-Seq experiment 2. C. UMAP plot of 128112 individual cellular transcriptomes highlighting the cell clusters. D. Bar diagrams showing the cell percentages in broad cell types and per cell types. E. UMAP plot of the neoblast score. F. Bar diagram showing the average neoblast score for each cell type.

We then sequenced these 10 sublibraries using Illumina sequencing and generated the gene expression matrices using our SPLiT-seq pipeline (see Methods). We integrated both experiments in one single-cell analysis using Harmony ^23^. We used Scrublet ^24^ to analyse potential doublets. We observed that several clusters had high percentage of doublets (Figure S1A-F) and removed cell barcodes with high doublet score. After these steps we obtained an integrated dataset with 128,112 cells in 91 cell clusters (Figure 1C, Figure S2, Figure S3A-D, Data S1-3). We grouped these clusters in broad cell type categories, including neoblasts, neurons, muscle, parenchymal (referred to as *cathepsin*+ cells in other single-cell studies ^25^), epidermis, phagocytes, secretory, protonephridia, pharynx, *psd*+ cells and basal/goblet cells (Figure 1D, Figure S3E). Further five clusters (containing 18-109 cells each, 211 cells in total, 0.16% of the total dataset) presented mixed profiles and were left unannotated.

To identify neoblasts based on their gene expression patterns, we generated a neoblast score from a set of neoblast markers, which we calculated for all cells (Figure 1E, Figure S4A-B) (see Methods). We then calculated the average neoblast score per cluster and observed that while neoblast clusters have the highest averages, clusters annotated as well characterised progenitors (e.g. early epidermal progenitors, neuronal progenitors) also have high neoblast scores. In addition, several secretory clusters also had relatively high average neoblast scores. We reasoned that unlike very abundant cell types such as the epidermis, where cells within different stages of differentiation are found in separated clusters, these relatively rarer cell clusters contained differentiated cell types together with their progenitors. We then used the average neoblast score to group clusters in three categories: neoblasts, clusters with progenitors, and differentiated cells (Figure 1F). Altogether, these analyses showed that our single-cell dataset had enough resolution to identify 91 cell clusters that included neoblasts, progenitor clusters, clusters with progenitor and differentiated cells, as well as clusters containing fully differentiated cells.

### 4C sorted cells show that mitotic neoblasts exhibit heterogeneous cell fate specialisation

To further characterise the clusters and their neoblast content, we analysed the distribution of neoblast scores in each cluster (Figure 2A), as well as the proportion of cells with high neoblast score (threshold 0.185) per cluster (Figure 2B). Most cells in neoblast clusters (1-9) had neoblast scores above the threshold (96.1%), indicating that the neoblast score, calculated from the expression of neoblasts markers, is a good classifier for neoblast identity. In contrast, most neuronal clusters except for the neuronal progenitors had scores below that threshold. This was similar in muscle, parenchymal clusters, and phagocyte clusters, with the progenitor cluster containing a higher proportion of cells with high neoblast scores than the respective differentiated cells (with the exception of epidermis 4). Interestingly, most secretory clusters, as well as the goblet cells, the *psd*+ cells and the basal cells, had higher proportions of cells with high neoblast score. While these clusters were relatively small, other comparably small clusters, such as the protonephridia, consisted mostly of cells with low neoblast scores. All clusters had cells with a wide range of neoblast scores. To have a better resolution and distinguish neoblasts, progenitors and differentiated cells within the clusters, we subdivided the clusters using MetaCell ^26^. This resulted in 6947 metacells with 18 cells each in average. For each metacell, we calculated the average neoblast score and classified them into metacells with high, medium and low neoblast score (thresholds: 0.258, 0.171) (Figure 2C). Neoblasts and progenitor clusters contain the largest proportion of metacells with high and medium neoblast scores, whereas differentiated clusters like neurons and muscle contain mostly metacells with low neoblast scores. Clusters annotated as differentiated cells and progenitors like the secretory clusters contain a mixture of metacells with high, medium and low neoblast scores. These results showed that many small clusters such as secretory, goblet and *psd*+ cells contain a notable proportion of cells that can be called neoblasts based on their high neoblast score, while many other small clusters such as neurons have very few.

**Figure 2:**
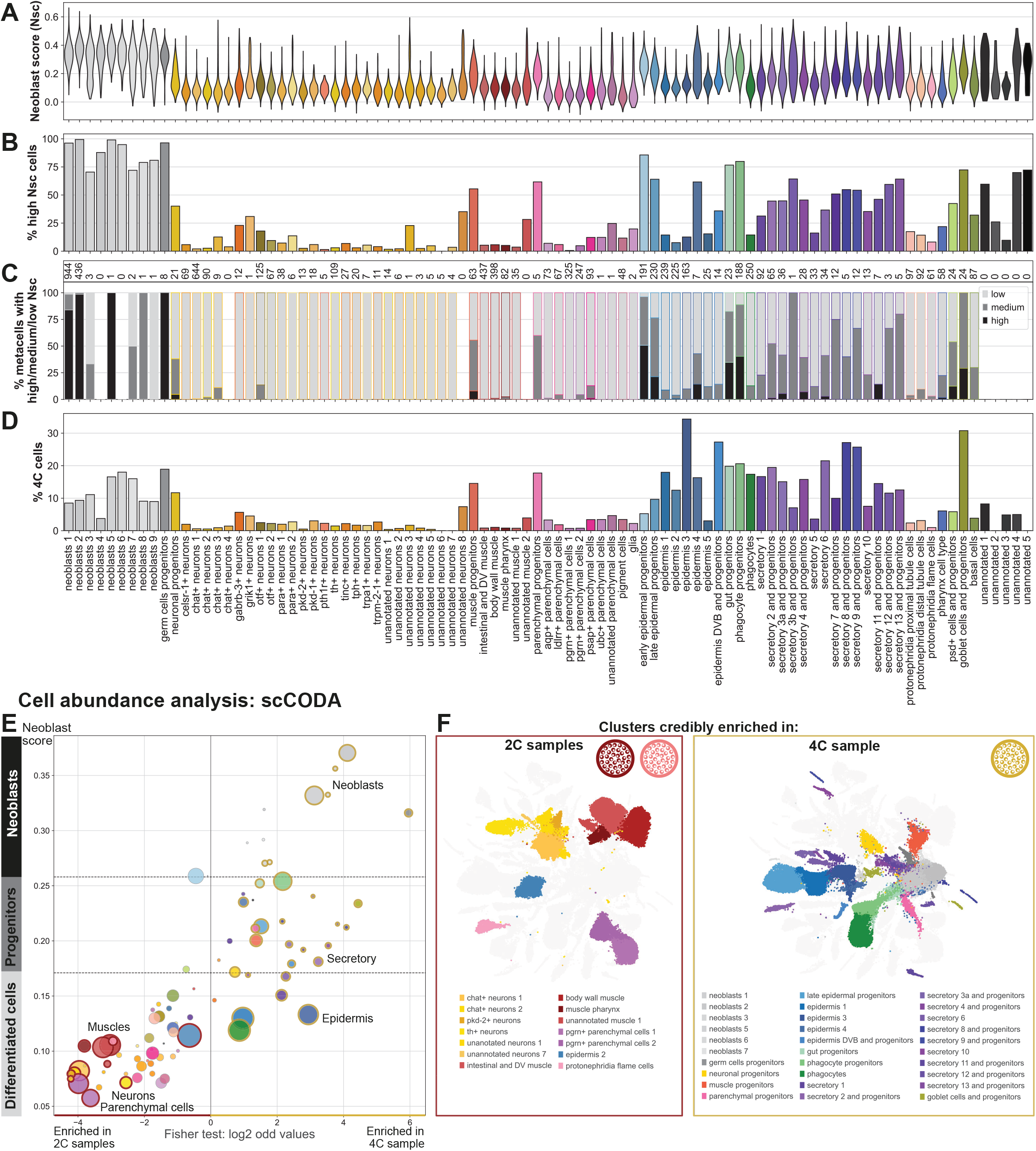
Neoblasts and progenitor clusters contain mitotic G2 neoblasts. A. Violin plots of the neoblast score per cluster. B. Bar diagram showing the percentage of high neoblast score cells per cluster. C. Metacell analysis of the dataset. The dataset was clustered using metacell and each metacell was labelled with the most frequent cell type label. The bar diagram shows for each cell type the percentage of metacells with a high, medium or low mean neoblast score (thresholds: 0.258 and 0.171). The number of metacells with the corresponding cell type label is indicated above. D. Bar diagram showing the percentage of 4C cells per cluster. E. Bubble plot of neoblast score against the log2 odd values calculated from a Fisher test comparing the distribution of 2C and 4C cells in each cell type. The size of the bubbles corresponds to the number of cells in the cluster and the colour to the cell types. The clusters with a golden and red borders are clusters defined as credibly enriched in 4C or 2C cells respectively according to the scCODA analysis (Fisher test, FDR= 0.05). F. UMAP plots highlighting the clusters credibly enriched in 2C cells and 4C cells according to the scCODA analysis.

We then wondered if those cells with high neoblast scores in progenitor clusters were mitotically active. Methods for cell-cycle inference based on reference gene sets assume either *a priori* knowledge of what genes are expressed during the cell cycle, or that cell cycle gene expression is conserved with that of orthologous genes from a reference model species. This assumption remains largely untested in planarians. Similarly, although *smedwi-1* and other markers used in the neoblast score are expressed in proliferating neoblasts, their levels decline as cells differentiate after cell cycle exit. Thus, it is not possible to determine mitotic activity from the neoblast score or cell-cycle scores alone. We resorted to our 4C sample (Figure 1A, Figure S3A), which contains cells sorted by their DNA content, duplicated after the S-phase of the cell cycle. Importantly, FACS sorting is an enrichment process that never results in a 100% efficiency: the 4C sample, even if highly enriched in mitotic cells, includes a small percentage of contaminant cells (here differentiated non-mitotic cells). Moreover, the 4C gate includes doublets, as they contain an amount of DNA equivalent to 4C cells. While we implemented a singlet gate as part of our routine FACS sorting procedure and further eliminate doublets bioinformatically using Scrublet ^23^, some doublets can still be present in the 4C sample. Thus, to rigorously identify clusters that contain mitotically active 4C cells we examined the relative enrichment of 4C gate cells by analysing the percentage of 4C gate cells in each cluster (Figure 2D). Consistent with the caveats explained above, all annotated cell clusters contained 4C cells. These proportions remained low (below 5%) in most differentiated cell clusters with the exception of epidermal cells in clusters epidermis 1 to 4, which were highly enriched in 4C cells. This may be attributed to a relatively high proportion of doublets in epidermal cells, likely arising from undissociated doublets tightly joint, or to the presence of multinucleated cells ^27^. All other clusters with high percentages of 4C cells were neoblast or progenitor clusters, including several small clusters of secretory cells, muscle, neuronal, phagocyte and goblet cell progenitors. Altogether these results indicated that mitotic neoblasts contained cells committed to several cell type fates. However, some fates are specified early, like for example the secretory cell types, but others, including neurons, muscle and phagocytes only acquire a general fate early, and diversify into the specific fates later in a post-mitotic phase of differentiation.

We then aimed at quantifying and statistically analysing the 4C content in each cluster. We checked whether the 2C and 4C samples were enriched in particular cell types using scCODA (Figure 2E-F). scCODA allows analysis of compositional changes of cell types between samples in single cell datasets thanks to a Bayesian model ^28^. The outcome of the scCODA analysis was consistent with the proportion of 4C cells highlighted above. The differential cell abundance analysis showed that differentiated cell types (notably neuronal, muscle and parenchymal cell types) were enriched in the 2C samples, whereas neoblasts and clusters containing progenitors, as well as epidermis, were enriched in the 4C sample. Notably, most of the clusters with a high neoblast score were enriched in the 4C sample, whereas clusters with a low neoblast score tended to be enriched in the 2C sample (Figure 2E). The exceptions were the epidermis and phagocytes enriched in the 4C sample. We also observed that 11/14 secretory cells clusters were enriched in the 4C sample. This analysis confirmed the fact that cells belonging to clusters with high neoblast score, i.e. neoblasts and progenitor clusters from all lineages, were enriched in the FACS sorted 4C sample. The enrichment in progenitor clusters indicated that those contain committed proliferating neoblasts.

### Neoblast fate commitment occurs sequentially during differentiation

To further ascertain the cell type fate identities present in 4C neoblasts, we performed several subclustering analyses. First, we randomly subsample cells from the 4C sample, computed the UMAP and performed a Leiden clustering using a procedure analogous to the one followed in our main original dataset (Figure 3A). To ascertain cell type fates, we applied 5 levels of Leiden clustering resolution, and identified the dominant original identity in each subcluster, representing at least 10% of the subcluster cells (Figure 3B). To assess the separation and robustness of the identified cell populations, we calculated average silhouette scores in each clustering ^29^ (Figure 3C). This subclustering analysis revealed that many individual secretory cell clusters were identifiable in the 4C sample. However, from broad type identities such as neurons and parenchymal cells, only the progenitor cluster as well as a few other cell types were present. For instance, only *otf*+ neurons 1 and 2 were identifiable at the lower resolution, and *celsr-1*+ and *para*+ neurons at higher resolutions. This indicated that these fates might diverge early from all other neuron fates and were observable among mitotic neoblasts. However, this analysis suggested that most other neuronal fates are acquired at a later post-mitotic stage of differentiation and were conversely undetectable in 4C samples. Similarly, from the parenchymal cell group, pigment, *aqp*+ and *psap*+ cells, as well as the parenchymal progenitors were identified, indicating that these fates diverge early in their differentiation process. From the different cell types that make the planarian gut, all types including gut phagocytes, goblet cells and basal cells were identified in 4C samples.

**Figure 3:**
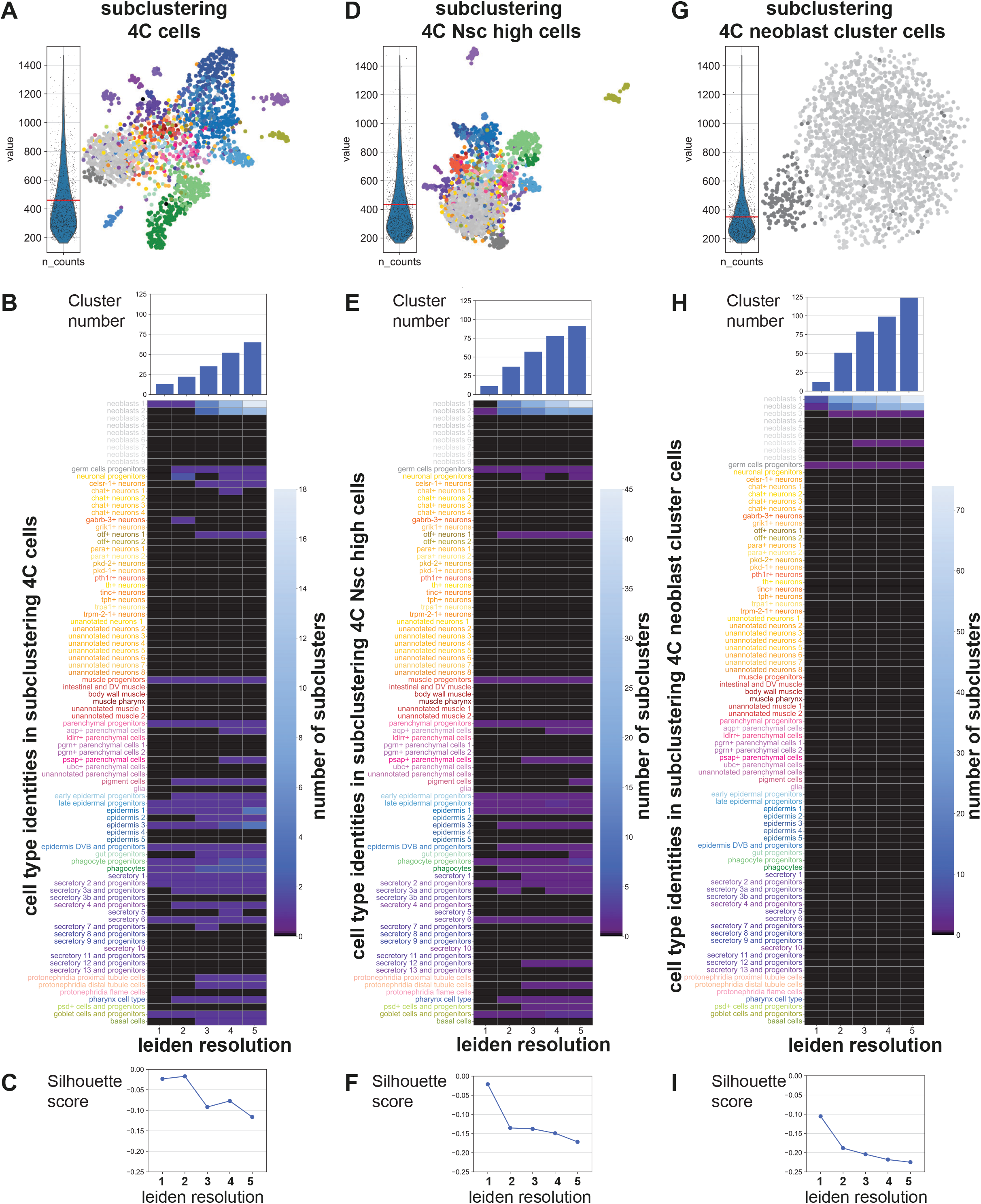
Subclustering of 4C cells reveals early fate commitment patterns in mitotic G2 neoblasts. A, D, G. Subclustering analyses of (A) all 4C cells, (D) 4C cells with a high neoblast score (>0.185), and (G) 4C cells annotated as neoblasts and germ cell progenitors. For 4C cells and 4C cells with high neoblast score, 1861 cells (i.e. the number of 4C cells annotated as neoblasts) were randomly subsampled. Each panel shows a violin plot of the number of UMIs per cell and a UMAP plot of the corresponding subset, with cells coloured according to their original cell type identity. B, E, H. Heatmaps showing, for the corresponding subclustering analyses in (A), (D), and (G), the number of clusters in which each original cell type label represents at least 10% of the cells. The barplot above shows the number of clusters for each clustering resolution. C, F, I. Dotplot showing the silhouette score for each clustering resolution of the 3 subclustering analysis.

We then wondered if doublets and contaminant cells could be driving those results. To further analyse this question, we performed a more stringent subclustering analysis, by randomly subsampling an equal amount of cells of the 4C sample with high neoblast scores (Figure 3D). We reasoned that contaminants should have low neoblast scores, corresponding to their differentiated cell type identity. This analysis confirmed that many cell type identities can be identified in this subset of cells. The individual identities largely correspond to those identified in the previous analysis (Figure 3E-F), lending further support to our findings.

Finally, to check whether further identities could be hidden within the neoblast clusters, we subset cells belonging to the neoblast broad category (i.e. neoblasts 1 to 9 and germ cell progenitors) and subjected them to an analogous processing pipeline (Figure 3G). Of note, only neoblasts and germ cell progenitor labels could be identified in this analysis (Figure 3H), as this is forced by the subset performed. The silhouette scores of the 4C neoblast subclustering condition is lower than the previous subsets (Figure 3C, F, I), revealing a lack of further structure in this dataset. This suggested that cells of the 4C sample in the broad category neoblasts correspond to unspecialised neoblasts. The UMI content of these cells was slightly lower than the previous sets (Figure 3A, D, G), however this minor difference is unlikely to explain the lack of differentiated cell fates in this subset.

Altogether, these analyses showed that many cell type fates could be identified in mitotic neoblasts. While several cell types—such as most secretory cell types—could be detected in several conditions, others, like neurons, were represented only by neuronal progenitors and a few early diverging subtypes. This indicated that neoblast cell fate commitment is sequential, and can happen early or late during their differentiation process. A large proportion of neoblasts did not show measurable commitment and likely correspond to unspecialised, naïve neoblasts.

### Gene expression characterisation of 2C and 4C cells

We then questioned if 2C and 4C uncommitted neoblasts had different gene expression patterns. We selected neoblasts as cells belonging to neoblast clusters with a high neoblast score (> 0.258), and performed differential gene expression analysis between the 2C and 4C samples using DESeq2. We identified 46 genes as differentially expressed. Among those, 18 were upregulated in 4C, 28 were upregulated in 2C neoblasts (Figure 4A, Figure S5, Data S4). Genes upregulated in 2C neoblasts were either expressed in most cell types or highly expressed in differentiated cells including early epidermal progenitors, phagocytes and parenchymal cells (Figure 4B-C, Figure S5A-B). On the other hand, most genes upregulated in 4C neoblasts were highly expressed in the neoblasts, with the exception of 3/18 being expressed in differentiated cells (Figure 4D-E, Figure S5C-D). Amongst these genes there were several histones (h1SMcG0006230, h1SMcG0008035, h1SMcG0009165), as well as *Smedwi-1* (h1SMcG0013999), a Tudor gene (h1SMcG0009920) and a Vasa homolog (h1SMcG0016303).

**Figure 4:**
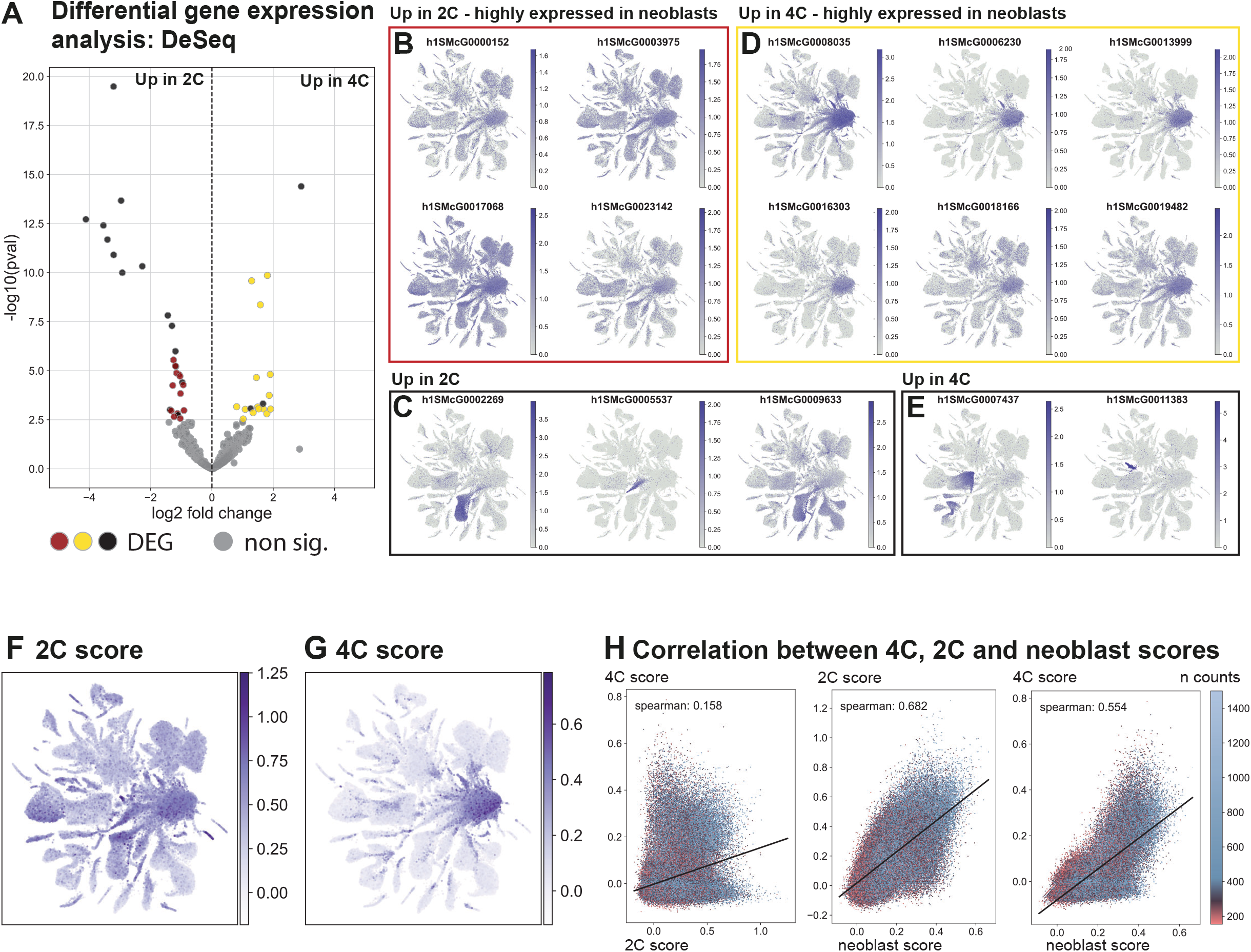
Differential gene expression analysis of 2C and 4C neoblasts. A. Volcano plot highlighting genes significantly upregulated in 2C neoblasts (red), genes significantly upregulated in 4C neoblasts (yellow), and significantly differentially expressed genes enriched in other cell types (black). B–E. UMAP feature plots showing the expression of representative differentially expressed genes from each category. F. UMAP plot showing the 2C score, calculated from the expression of genes upregulated in 2C neoblasts. G. UMAP plot showing the 4C score, calculated from the expression of genes upregulated in 4C neoblasts. H. Correlation between the neoblast score and the 2C and 4C scores. Cells are coloured according to their UMI counts to highlight low- and high-quality cells (red and blue, respectively).

Importantly, we did not observe cell cycle genes such as cyclins and CDKs populating this analysis, suggesting that their expression might not be the dominant component of the transcriptomic profile of planarian neoblasts. While this feature is often taken as an assumption, it is possible that cell cycle specific gene transcription results in transcripts that are detectable throughout the cycle and therefore are not detected in our analysis. Alternatively, the experimental design with only two conditions and sparse SPLiT-Seq data, may have lacked the sensitivity to detect finer grain RNA level oscillations.

The differential gene expression analysis identified gene sets enriched in 2C and 4C uncommitted neoblasts. We used those sets to generate 2C and 4C scores respectively (Figure 4F-G). Consistent with the expression patterns of the individual genes, the 2C score was higher in the neoblasts and progenitors and showed medium values in the differentiated cell types. In contrast, the 4C score was the highest in the neoblasts and in progenitor clusters, and had very low values in the differentiated cells. The two gene sets were non-overlapping and had distinct expression profiles resulting in an absence of correlation between the 2C and the 4C scores (Figure 4H). On the contrary, there was a medium correlation between the 2C and 4C scores and the neoblast score (Figure 1E). The correlation can be explained partly by genes present in both the neoblast score and the 2C or 4C scores (respectively 7 / 13 genes common for the neoblast score and the 2C score, 12/15 genes common for the neoblast score and the 4C score). The gene set used to generate the neoblast score contained an additional 39 genes that were not seen in this analysis and were likely not linked to cell cycle phases.

### Simplified regulation of cell cycle exit in *Schmidtea mediterranea*

Planarians are known to have lost key cell cycle genes compared to vertebrates. A few studies have reported the presence and absence of cell cycle genes in *Schmidtea mediterranea* using previous versions of the genome as reference ^18,30,31^. To characterise cell cycle regulators in our dataset, we first used OrthoFinder ^32^ and a set of model organism species with cell cycle annotation to look for orthologs in our reference genome ^33^ (Data S5, Data S6, Data S7, Figure 5A). 1175 genes were identified, including key cell cycle regulators like 4 cyclins, 5 aurora kinases and members of the APC complex. Our analysis retrieved a binding partner of the APC complex, *cdh1* (annotated as Cdc20 due to their sequence similarity, we confirmed the identity using the *Smed-cdh1* sequence from Sato *et al*). In *Dugesia japonica*, *cdh1* is responsible for cell cycle exit ^19^. We identified Cdc20, the other binding partner, with a reciprocal blast using the *Smed-cdc20* sequence from Sato *et al*. Similar to *Dugesia*, we could not find regulators of *cdh1* and CDK inhibitors. Indeed, OrthoFinder did not retrieve any sequence for CDK inbitors (INK family: p15, p16, p18, p19, CIP/KIP family: p21, p27, p67) and the regulators of APC/C-*cdh1* Emi1 and SCF-Skp2. Our analysis supported the hypothesis, that similarly to *Dugesia japonica*, *cdh1* could be the only point of control regulating cell cycle exit and differentiation of the neoblasts (Figure 5A).

**Figure 5:**
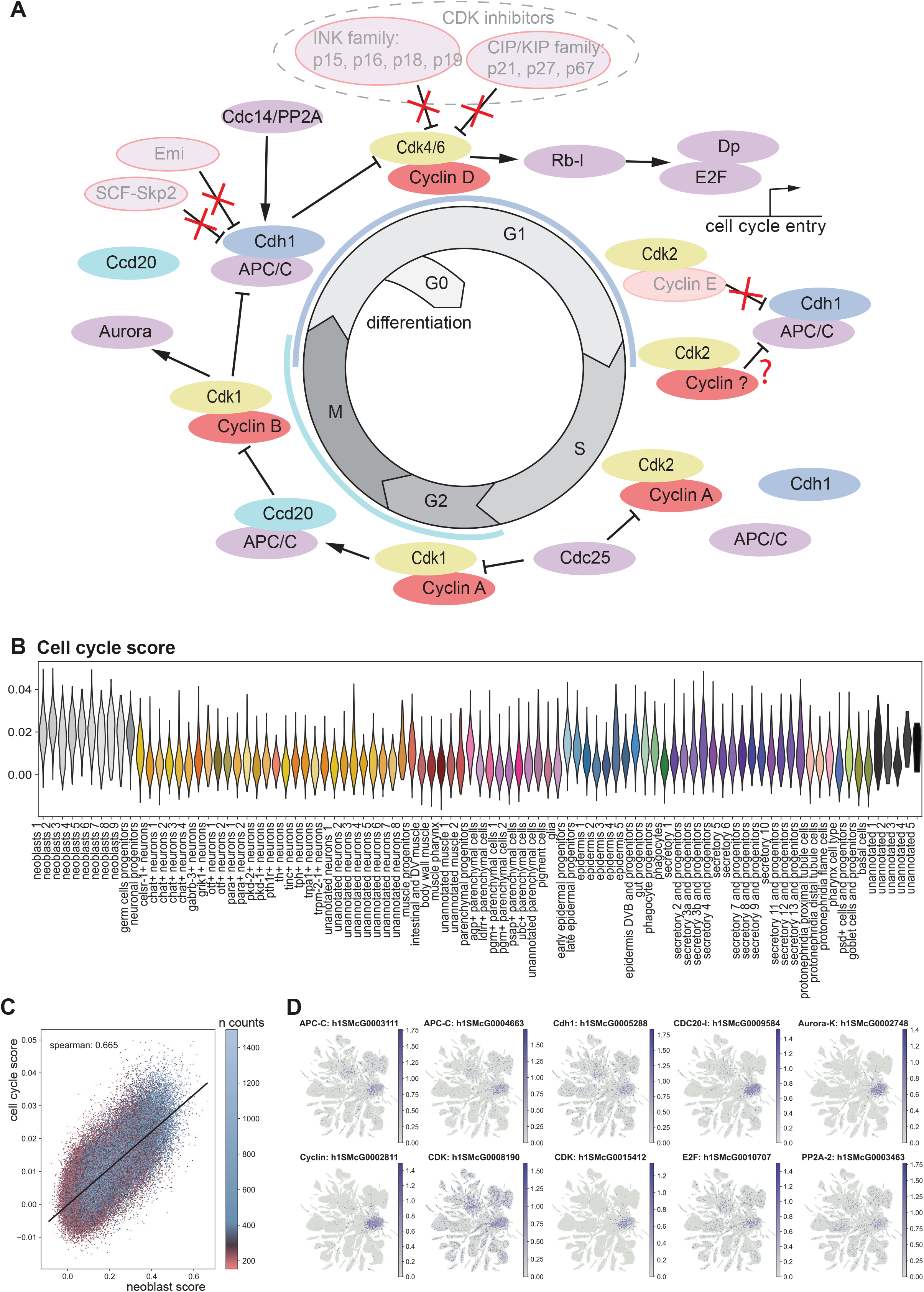
Characterisation of cell cycle genes in *Schmidtea*. A. Schematic of the main actors of the cell cycle regulation identified with OrthoFinder. The genes with the red outline were not found and we assume those genes to be missing in *Schmidtea*. B. Violin plot of the cell cycle score. C. Correlation between the cell cycle score and the neoblast score. The cells are coloured according to their UMI numbers to highlight low and high quality cells, respectively in red and blue. D. UMAP feature plots showing the expression of example cell cycle genes. The genes names were determined using OrthoFinder.

Then we checked the clusters in which those genes are expressed. Using differential gene expression analysis between the cells belonging to the neoblast clusters and all other cells, we determined that 448 genes have an enriched expression in the neoblasts (Data S8). A cell cycle score generated with those 448 genes followed the same tendency as the neoblast score: it was the highest in neoblast and germ cells progenitors, and higher in clusters containing progenitors than differentiated cells (Figure 5B, Data S8). 10 genes were common between the cell cycle score and the neoblast score, and the Spearman coefficient showed positive correlation between those scores (Figure 5C). The genes identified as key cell cycle regulators were expressed predominantly in the neoblasts, consistent with a role in cell cycle regulation (Figure 5D, Figure S6). Altogether, our analysis supported the gene loss observed in *Schmidtea* and highlighted the potential key role of *cdh1* in cell cycle regulation.

### Knockdown of *cdh1* and *h2b* impact homeostasis and regeneration

*cdh1* was characterised as a key regulator of cell cycle exit in *Dugesia* ^19^. However, neoblast populations, specialisation and heterogeneity are much better understood in *Schmidtea*. To integrate these two subjects and to investigate the role of *cdh1* in the neoblasts of *Schmidtea* at the single-cell level, we performed knockdown by RNA interference followed by scRNA-seq. Since *cdh1* RNAi could lead to hyperproliferation of the neoblasts, we added *h2b* RNAi known to deplete neoblasts and progenitors as opposite effect ^22^.

dsRNA was incorporated in food pellets and the animals were fed 5 times over 2 weeks (Figure 1B). 17days after the 1^st^ feeding, we scored the phenotypes and dissociated animals with ACME (Figure 6A). All animals fed with dsRNA for *gfp* presented entirely intact bodies without any phenotype visible under the stereomicroscope. From the animals fed with dsRNA for *cdh1*, 63 did not have any visible phenotype, 24 had dark lesions on the body, 12 died during the feeding. Animals fed with *h2b* dsRNA either died (n=6), presented dark lesions on the body (n=14), had head regression (n=15) or no phenotypes (n=64). 13 animals from each condition and replicate were cut transversally in the middle of the pharynx, making a head fragment without tail and a tail fragment without head.

**Figure 6:**
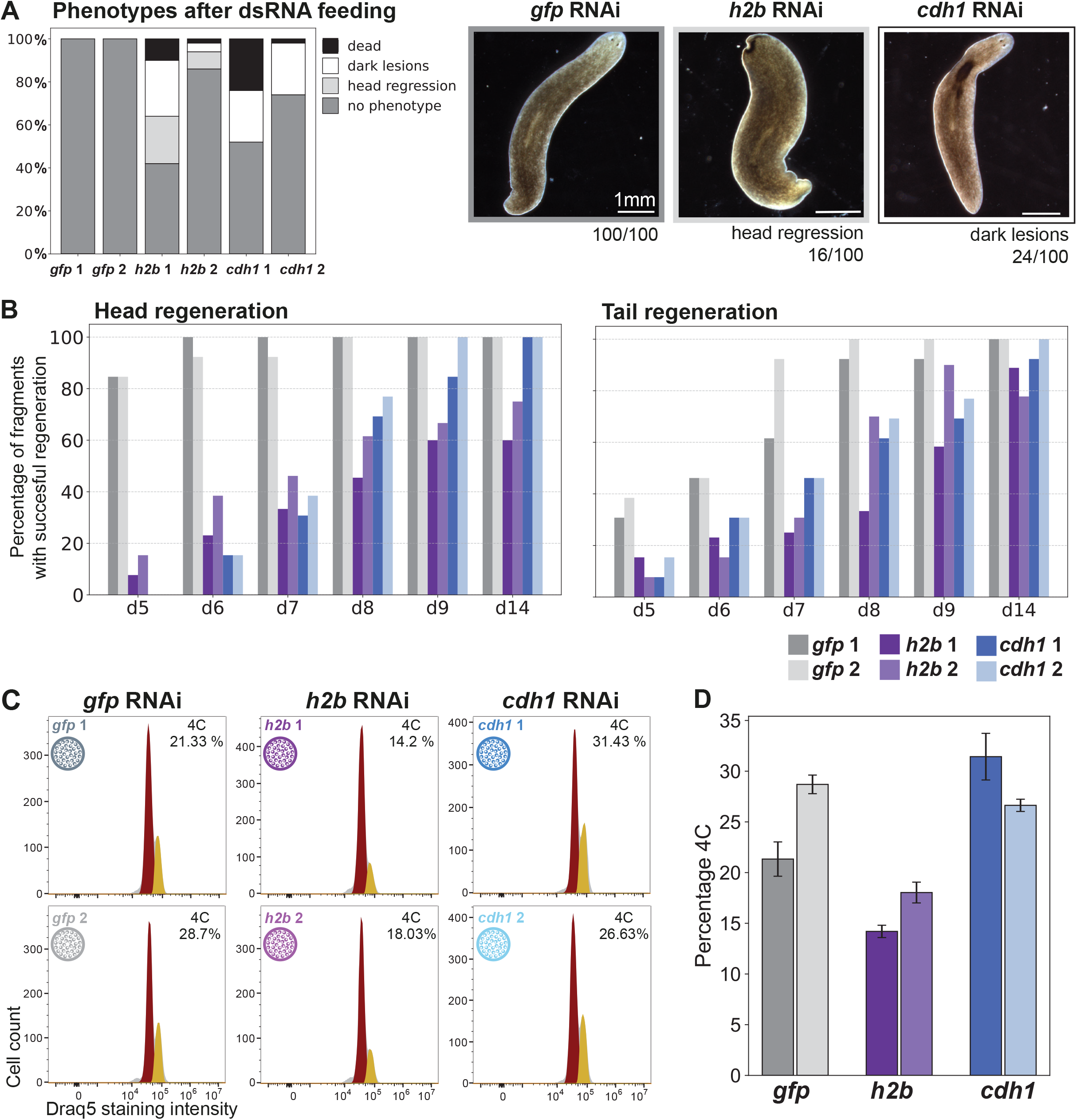
Knockdown of *cdh1* and *h2b* impairs homeostasis and regeneration. A. Phenotypes of intact animals after feeding 5 times with dsRNA over 17 days. B. Head and tail regeneration after transversal section for the 3 dsRNA feeding conditions (*gfp*, *cdh1*, *h2b*). Results are in percentages of the living animals. In the head regenerating fragments of the *h2b* RNAi, 4 animals died (3 for replicate 1 and 1 for replicate 2). In the tail regenerating fragments of the *h2b* RNAi, 8 animals died (4 per replicate) as well as 1 fragment of *cdh1* replicate 2. C. Cytometry profiles of cell dissociations of entire animals after dsRNA feeding and quantification of the 4C population. D. Barplot showing the percentage of 4C cells in each sample. Each sample was measured 3 times, the bars show the average of the 3 measurements.

We scored regeneration of the head and tail and considered regeneration as complete when respectively two eyes or the pointy tail were visible (Figure 6B). Head and tail regeneration proceeded rapidly in control animals but was substantially delayed following *cdh1* knockdown and strongly impaired following *h2b* knockdown. While nearly all *cdh1* RNAi animals eventually regenerated completely with a delay of 6 days compared to control animals, only 60-80% of the *h2b* RNAi animals regenerated completely by the end of the experiment. These experiments confirmed that knockdown of *cdh1* by dsRNA feeding leads to dark lesions in the body of intact animals and death of some animals, as well as a delay in head and tail regeneration, similar to the phenotypes observed in *Dugesia japonica* ^19^. Knockdown of *h2b* led to head regression, lesions in the body and death of intact animals and strongly impaired regeneration. Those phenotypes are milder, likely due to the dsRNA delivery by feeding, but consistent with those reported by Solana and coworkers.

### *cdh1* and *h2b* knockdown have opposite effects on the 4C cell population

*cdh1* and *h2b* RNAi show similar phenotypes on the intact worms with the presence of lesion and death of some animals. In both cases, regeneration is also impaired. However, their impact on the neoblast population is opposite: *h2b* is required for neoblast maintenance ^22^, but *cdh1* is responsible for the cell cycle exit of the neoblasts ^19^. To characterize cell population and gene expression changes after *cdh1* knockdown, we first assessed the proportions of 2C and 4C cells by cytometry after ACME dissociation, using the fluorescence intensity of the nuclear DRAQ5 staining (Figure 6C, D). For the animals fed with dsRNA for *gfp*, the proportion of G2 was between 21% and 28% but for animals fed with *cdh1* dsRNA the proportion was higher (26-31%). This indicates that cells are either arrested in the G2 phase, or that more cells are entering the cell cycle compared to the control condition. The proportion of G2 was lower (14-18%) for the animals fed with *h2b* dsRNA, consistent with a depletion of the neoblast populations ^22^.

To characterize cell populations at the single-cell level, we generated a single cell RNA-seq dataset after completing the RNAi procedure (Figure 1B, Figure S3A). As a comparison, we included animals fed with *h2b* dsRNA and controls fed with *gfp* dsRNA. We analysed the cell proportions of each cluster using scCODA ^28^. This analysis revealed distinct and opposing effects of *cdh1* and *h2b* RNAi on the neoblast clusters (Figure 7A, Figure S3A). Following *cdh1* RNAi, the 6 largest neoblasts clusters are significantly enriched. The two largest clusters represent together 90.5% of the total number of neoblasts and 12.9% of the total number of cells in control conditions. After *cdh1* RNAi, they represent 28.0% of the total cell number. In the case of *h2b* RNAi, the 2 largest neoblast clusters are significantly depleted, representing 5.77% of the total number of cells. This is consistent with the respective increase and decrease of the 4C cell population observed by cytometry. The increase of the neoblast clusters observed on the single cell RNA-seq data following *cdh1* RNA is higher than the increase of 4C cells observed by cytometry suggesting a global increase of the neoblast population and not only an increase of 4C cells.

**Figure 7:**
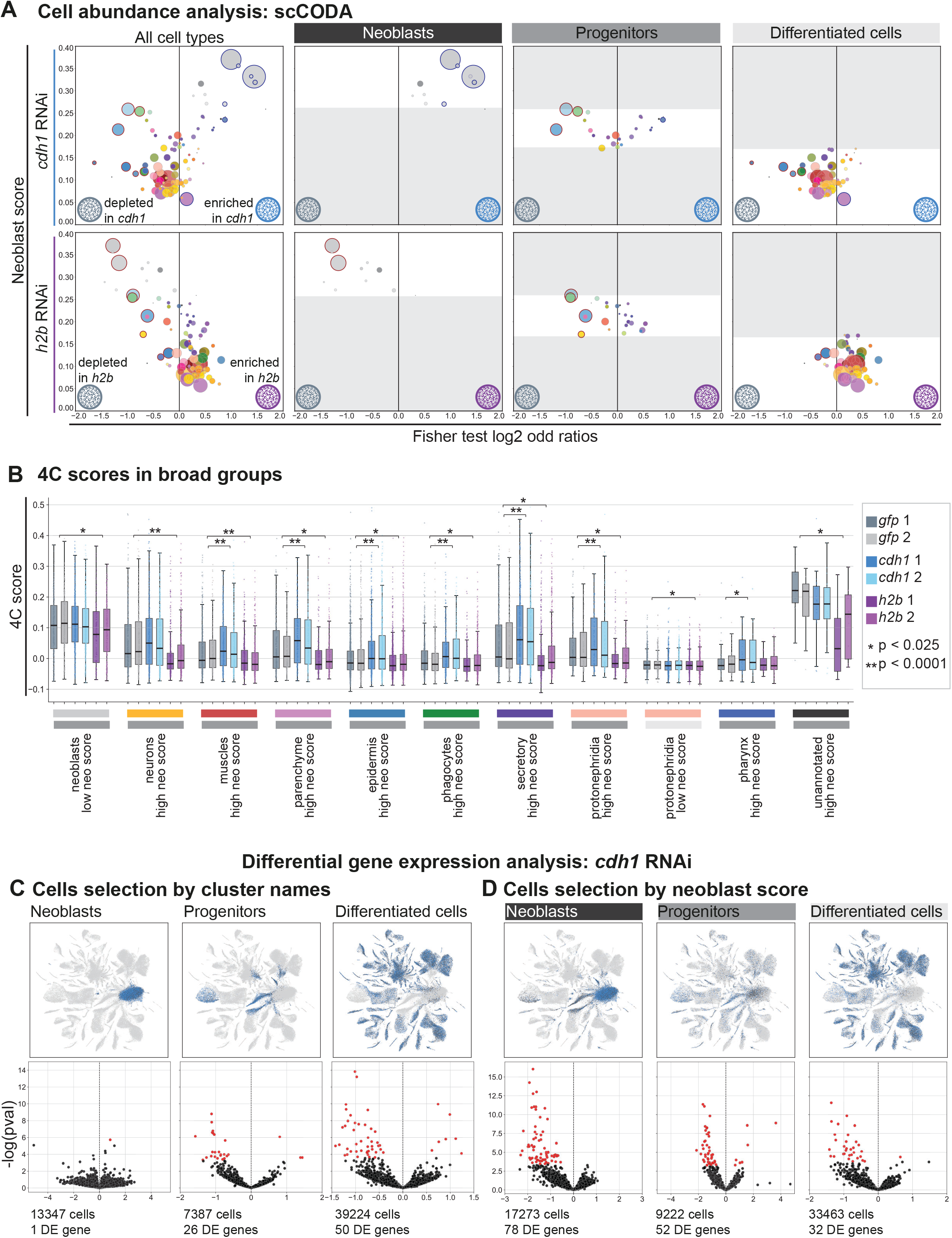
*cdh1* affects both neoblasts and progenitors. A. Bubble plot showing the neoblast score versus the log2 odds ratio from a Fisher’s exact test comparing the distribution of *cdh1* or *h2b* RNAi cells with *gfp* RNAi controls across cell types. Bubble size indicates the number of cells, and colour indicates the cell type. Blue- and red-outlined bubbles denote clusters identified as credibly depleted or enriched, respectively, in *cdh1* or *h2b* RNAi relative to *gfp* RNAi by scCODA (FDR = 0.05). B. Box plots showing the 4C score for each RNAi treatment across broad cell type categories. Only the categories with significant differences are presented here. The categories without significant differences are shown in Figure S7. Cell types were further classified into high- and low-neoblast-score groups based on their neoblast score. The asterisks indicate significant differences from the *gfp* RNAi control according to a linear mixed-effects model. Only significant comparisons are shown. C. Top: UMAP feature plots showing the cells selected for differential expression analysis, grouped into neoblasts, progenitors, and differentiated cells according to cluster identity. Bottom: Volcano plots highlighting in red genes significantly up- or downregulated in *cdh1* RNAi relative to *gfp* RNAi. D. Top: UMAP feature plots showing the cells selected for differential expression analysis, grouped into neoblasts, progenitors, and differentiated cells based on their neoblast score. Bottom: Volcano plots highlighting in red genes significantly up- or downregulated in *cdh1* RNAi relative to *gfp* RNAi.

### Cell cycle exit is essential for differentiation but not for neoblast specialisation

We next assessed the effects of *cdh1* and *h2b* RNAi on the remaining cell populations (Figure 7A). Interestingly, the effects of *cdh1* and *h2b* RNAi on progenitor populations were not opposite; instead, both treatments resulted in depletion of progenitor clusters. Epidermal progenitors (early and late) accounted for 8.53% of cells in the *gfp* RNAi control, but only 4.13% and 5.14% following *cdh1* and *h2b* RNAi, respectively. Similarly, phagocyte progenitors decreased from 2.29% in the control to 1.54% and 1.40% after *cdh1* and *h2b* RNAi, respectively. Neuronal progenitors were also reduced following *h2b* RNAi. These results suggest that post-mitotic progenitors become depleted in both conditions as differentiating cells fail to be replenished by the neoblast population. Following *h2b* RNAi, this depletion is explained by the loss of neoblasts, as previously reported ^22^. In contrast, in *cdh1* RNAi, the reduction of the progenitor populations can be explained by the accumulation of neoblasts that fail to exit the cell cycle and to generate postmitotic progenitors. Differentiated cell types were only modestly affected by either treatment (Figure 7A). Following *cdh1* RNAi, epidermal and phagocyte populations were reduced, whereas a parenchymal cluster and one epidermal cluster were enriched, suggesting lineage-specific sensitivities to disruption of cell-cycle regulation. In contrast, *h2b* RNAi resulted in enrichment of pharyngeal cells.

To investigate how *cdh1* affects cell cycle progression, we used the 2C and 4C cell cycle scores derived from genes differentially expressed between 2C and 4C cells experiment (Figure 4F-G). These scores provide a proxy for cell cycle state, with a high 4C score indicating enrichment for G2/M-associated cells. To assess the effects of *cdh1* and *h2b* RNAi across cell types, we quantified the 2C and 4C scores within each broad cell class. Each broad cell type category was further subdivided into cells with high neoblast scores (higher than 0.171), representing cycling specialised neoblasts, and cells with low neoblast scores, representing differentiated cells (Figure 7B). Likewise, neoblasts were separated into cells with high neoblast scores (higher than 0.258) and low neoblast scores.

Following *h2b* RNAi, the 4C score was significantly reduced in neoblasts with low neoblast scores and in the high-neoblast-score populations of most broad cell types, including neurons, muscle, parenchyma, epidermis, phagocytes, secretory cells, protonephridia and unannotated cells. This reduction is consistent with the known depletion of proliferating cells following *h2b* RNAi. In contrast, *cdh1* RNAi increased the 4C score in the high-neoblast-score populations of muscle, parenchyma, epidermis, phagocytes, secretory cells, protonephridia and pharynx. This high neoblast score population is very likely to be composed of specialised neoblasts that are categorised as differentiated cells because of the expression of FSTFs. The increase of the 4C score following *cdh1* RNAi indicates that similarly to uncommitted neoblasts, this population does not exit the cell cycle. Contrary to *h2b* RNAi, the neoblast broad group was not affected by *cdh1* RNAi. These opposing effects between *cdh1* and *h2b* RNAi suggest that, whereas *h2b* RNAi depletes proliferating progenitors, *cdh1* RNAi leads to the accumulation of cycling specialised neoblasts that fail to exit the cell cycle, accompanied by a corresponding depletion of postmitotic progenitors.

Neoblasts with a low neoblast score, as well as differentiated cells with a low neoblast score did not show any significant changes after *cdh1* and *h2b* RNAi (Figure S7A), indicating that fully differentiated cells are less sensitive to these perturbations of cell cycle regulation. The 2C score was less affected overall. We nevertheless observed an increase in muscle clusters with high neoblast score following both *cdh1* and *h2b* RNAi, as well as increases in phagocytes and secretory progenitors after *cdh1* RNAi and a decrease in unannotated cells after *cdh1* RNAi (Figure S7B-C). Together, the pronounced effects on the 4C score and the limited changes in the 2C score indicate that the actively cycling cells are the primary targets of RNAi-induced perturbations.

Finally, to understand the genetic basis of these cellular changes, we examined gene expression following *cdh1* RNAi in neoblasts, progenitors, and differentiated cells. Differential gene expression analysis was performed using DESeq2 on pseudobulk samples generated by aggregating single-cell transcriptomes within each population (Figure 7C-D).

In a first analysis, cells were grouped according to their broad cell type annotation into neoblasts, progenitors, or differentiated cells. Under this classification, neoblasts were largely unaffected, with only a single differentially expressed gene, whereas progenitors and differentiated cells showed 26 and 50 differentially expressed genes, respectively (Figure 7C, Data S9). These results indicate that the transcriptional state of unspecialised neoblasts is largely unaffected by *cdh1* knockdown.

We next repeated the analysis after subdividing cells according to their neoblast score, allowing specialised neoblasts to be included within the neoblast population. This approach identified 78 significantly differentially expressed genes in neoblasts, 52 in progenitors, and 32 in differentiated cells (Figure 7D; Data S10). Most of these genes were downregulated, and many were normally expressed in progenitor populations, including early and late epidermal progenitors, phagocyte progenitors, and differentiated cell types. These results suggest that specialised neoblasts fail to activate transcriptional programs associated with differentiation following *cdh1* RNAi, despite retaining their specialised identity.

Notably, the genes affected by *cdh1* RNAi were largely distinct from those differentially expressed following *h2b* RNAi (Figure S8 A-B), indicating that although both perturbations reduce progenitor populations, they do so through mechanistically distinct pathways. Together, these findings support a model in which specialised neoblasts fail to exit the cell cycle and initiate differentiation, consistent with the accumulation of cells with high 4C scores. Our single-cell analysis therefore reveals that cell cycle exit is required for differentiation but not for neoblast specialisation.

## Discussion

Investigation of cell cycle and differentiation in *Schmidtea* has relied so far mainly on experimental procedures that do not reflect physiological conditions. Our aim was to expand that knowledge by studying homeostasis conditions in fixed samples. To investigate the relationship between cell cycle regulation and differentiation in *Schmidtea*, we generated two scRNA-seq datasets from uninjured animals, including libraries enriched for 2C and 4C DNA content and libraries generated after knockdown of cell cycle regulators. Our analysis revealed that progenitor clusters across multiple lineages contain actively cycling cells enriched in the 4C state, indicating that fate commitment occurs before completion of the cell cycle. Orthology analysis highlighted a simplified repertoire of cell cycle regulators in *Schmidtea*, suggesting a central role of *cdh1* in controlling cell cycle exit. We further investigated the role of *cdh1*, using RNA interference of *cdh1* and *h2b*. While both perturbations lead to homeostasis defects and impaired regeneration, they had opposite effects on the cycling cell population, with *cdh1* RNAi leading to accumulation of 4C cells and neoblast enrichment. These results support a model in which lineage commitment is initiated in cycling neoblasts, whereas terminal differentiation depends on *cdh1*-mediated cell cycle exit.

Our 4C sample enriched in cells in G2 phase allowed us to characterise proliferating neoblasts and their progeny. Identifying those in single cell datasets is complex because cell cycle regulation in planarians has not been fully described yet. Moreover, cell cycle regulation consists of a reduced set of components compared with other metazoans, so models developed for vertebrates might not work as intended. In previous scRNA-seq datasets, neoblasts were identified using the expression of *Smedwi-1* ^15,16,20,25,34,35^. However, single cell datasets are sparse so relying on a single gene may fail to capture all relevant cells. Therefore, we implemented a neoblast score based on the 50 top genes of the two neoblast clusters of our analysis to identify cells with neoblast identity. Cells with a high neoblast score were enriched in the 4C sample, validating our approach. This neoblast score also provides a way to distinguish specialised neoblasts and postmitotic progenitors. Indeed, neoblasts and other cells with a high neoblast score accumulate following *cdh1* RNAi and are depleted following *h2b* RNAI. This behaviour is consistent with an identity as neoblasts. On the contrary, cells with a low neoblast score are mildly affected by those RNAi treatments, indicating an identity as postmitotic cells.

Analysis of the 4C sample alone revealed a high heterogeneity of the cycling cells in homeostasis conditions. Many cells have a gene expression profile that allows to assign them to a differentiated cell cluster, indicating that commitment can occur while cells are still in S/G2/M phases. This is seen when taking all cells, also when subclustering only progenitors using the neoblast score. This observation is consistent with previous reports highlighting expression of fate-specific transcription factors (FSTFs) in proliferating neoblasts ^14,36^.

Comparative analyses indicate that planarians possess a simplified cell cycle regulatory network in which *cdh1* plays a central role in promoting cell cycle exit and differentiation. Our findings in *Schmidtea mediterranea* largely agree with previous work in *Dugesia japonica*, supporting a conserved function for *cdh1*. Together, these results suggest that the role of *cdh1* as a key regulator of cell cycle exit arose early in planarian evolution and has been maintained despite extensive loss of other cell cycle regulators, highlighting planarian neoblasts as a valuable model for studying the coupling of proliferation and differentiation in a simplified regulatory network.

Knockdown of *cdh1* by RNAi leads to the accumulation of 4C neoblasts, resulting in defects in homeostasis and regeneration. While lineage commitment appears to begin before mitotic exit, neoblasts must subsequently withdraw from the cell cycle to complete differentiation. This model is consistent with previous studies in *Dugesia japonica*, in which knockdown of *cdh1* by RNAi leads to hyperproliferation of the neoblast and decrease of progenitors ^19^. Our analysis further dissects the boundary of that decision. Cells can initiate commitment while cycling, but successful differentiation requires cell cycle exit. As a result, after *cdh1* inhibition, mitotic unspecialised or specialised neoblasts accumulate, while post-mitotic progenitors become scarce.

It has been shown that neoblasts express FSTFs preferentially during the G2/M phases of the cell cycle ^3^. Following this model, neoblasts under *cdh1* RNAi may either fail to express FSTFs, or express FSTFs and not differentiate due to cell cycle related mechanisms. In our dataset we could not see any evidence for differential expression of FSTFs in *cdh1* neoblasts. This seems to support the fact that neoblasts are still able to commit to different fates despite *cdh1* knockdown. However, the absence of difference could also be due to the sparsity of SPLiT-Seq data.

King *et al* ^14^ suggested that among the 125 cell types/states of planarians, specification occurs at different stages. According to this framework, muscle, epidermal, parenchymal, and intestinal identities emerge within mitotic progenitors, whereas neuronal diversity increases at the post-mitotic progenitor stage, and protonephridial and phagocyte identities are established only during terminal maturation. This analysis was done with regenerating worms. Our analyses in uninjured worms broadly support this hierarchical model. Subclustering of 4C cells shows a diversity of transcription profiles in the mitotic cells, particularly in the gut cell types, secretory cells, and epidermis. This confirms King *et al* conclusion about the early specification in epidermis, parenchyma and intestinal cells and shows that this process happens in physiological conditions. Contrary to this study, we could not identify muscle types in 4C cells. We could also see some neuronal cell types in 4C cells, in higher number when taking all cells compared to selecting only 4C cells with a high neoblast score, strengthening the model of diversification at the progenitor level.

Our results resolve an important distinction between neoblast specialisation and subsequent differentiation. King *et al*. established that neoblasts can acquire lineage-specific transcriptional signatures while still in the cycling S/G2/M state ^14^, but whether these fate choices require cell-cycle exit remained unknown. Conversely, Sato *et al*. showed that *cdh1* knockdown causes neoblast accumulation and depletion of progenitors and differentiated cells, supporting a requirement for cell-cycle exit for differentiation ^19^, but did not determine whether neoblast specialisation itself depends on exiting the cell cycle. By combining cell-cycle-resolved single-cell profiling with functional inhibition of *cdh1*, we distinguish these two processes. In *cdh1* knockdown animals, specialised neoblast populations remain detectable at comparable levels, indicating that specialisation occurs in actively cycling neoblasts and therefore does not require cell-cycle exit. In contrast, the subsequent progression of these cells into progenitors and their diversification into differentiated cell types is strongly impaired. Thus, our results place cell-cycle exit after neoblast specialisation but before terminal differentiation and fate diversification, revealing distinct cell-cycle requirements at successive stages of planarian cell fate acquisition.

Thus, our results support a model in which lineage commitment is initiated during the cell cycle. The full diversity of cell types emerges at different stages depending on the lineage. This is strengthened by the fact that progenitors are present in the 4C sample, indicating that cells classified as progenitors are cycling cells. Cell cycle progression and differentiation are therefore tightly coordinated processes rather than sequential events, with neoblasts progressively acquiring differentiated identities before exiting the cell cycle.

## Conclusion

Our study demonstrates that neoblast specialisation and differentiation are distinct sequential processes. While many mitotic neoblasts are already committed to specific cell fates, further commitment occurs at different stages depending on the lineage, leaving a substantial pool of unspecialised neoblasts. Evolutionary analyses support a conserved role for *cdh1* in cell cycle regulation, and functional experiments reveal that *cdh1* is dispensable for neoblast specialisation but essential for cell cycle exit and the activation of differentiation programs. Consequently, *cdh1* depletion leads to the accumulation of cycling specialised neoblasts and the loss of post-mitotic progenitors, establishing cell cycle exit as a prerequisite for differentiation rather than for cell fate specification.

### Limitations of the study

This study has several limitations that should be considered. First, SPLiT-seq datasets are sparser than those generated using commercial droplet-based scRNA-seq methods, which may reduce the detection of lowly expressed cell cycle regulators and contribute to differences from previous single-cell atlases. The increased sparsity can also affect the performance of bioinformatic approaches, including cell cycle phase assignment based on gene expression. Second, enrichment of the 4C population by FACS is not perfect, and occasional doublets or cell aggregates may account for a small fraction of differentiated cells detected in this population. Nevertheless, the strong enrichment of neoblasts and progenitors supports our conclusions. Finally, although our results identify *cdh1* as a central regulator of cell cycle exit, we cannot exclude indirect effects of *cdh1* RNAi on other components of the cell cycle machinery. Future studies will be required to distinguish the direct and indirect consequences of *cdh1* depletion.

## Materials and Methods

### Animal culture

The worms used for the experiments were asexual *Schmidtea mediterranea*, derived from the clonal line Berlin-1 ^37^. They were kept at 18-20C in 1x Montjuic water (1.6 mM NaCl,1.0 mM CaCl2, 1.0 mM MgSO4, 0.1 mM MgCl, 0.1 mM KCl, and 1.2 mM NaHCO3, dissolved in deionised water) at pH 7.0. They were fed with frozen cow liver once to twice per week and starved for one week before any experimental procedure. All animals collected for the experiments were between 5-10 mm.

### ACME dissociation and cytometry analysis of the cell suspensions

Dissociation and fixation of the animals was performed using ACME ^20^ as described in Pérez Posada *et al* ^15^. Dissociated cells were stored in 1% PBS-BSA with 10% DMSO at −80C. Prior to library preparation, one aliquot of each sample was analysed by cytometry (CytoFlex S Flow Cytometer, Beckman Coulter). Samples were washed with 1% PBS-BSA, filtered with a 100um strainer (CellTricks yellow, Wolfslab) and stained with DRAQ5 and concanavalinA-FITC (1mg/mL stock diluted 1:250 in sample). We used the gating strategy described in Garcia Castro *et al* ^20^ to assess using the percentage of single cells and the percentages of G1/G2 populations, and hence evaluate the quality of the samples before library preparation. FlowJo v10 was used to quantify the 2C and 4C cell populations based on DNA content as determined by DRAQ5 staining, using a similar gating strategy.

### Gene knockdown by RNA interference (RNAi)

**cDNA synthesis:** RNA was extracted from wildtype *S. mediterranea* worms with TRIzol (Invotrogen) following the manufacturer’s protocol, and was used as template for cDNA synthesis using maxima H minus Reverse transcriptase (ThermoFisher scientific).

**Amplification of target genes:** cDNA was used as template for the amplification of *cdh1* and *h2b*. The primer sequences were: ggccgcggATTTACGCGACACCAGAACC (*cdh1*-F), gccccggccTACACGAGAGCGACTGATGG (*cdh1*-R), ggccgcggGCTGTTACAAAATACACAGGA (*h2b*-F) and gccccggccTCCTGTGTATTTTGTAACAGC (*h2b*-R). The primers for *cdh1* were designed based on the blastn best hit for the *cdh1* sequence from *Dugesia japonica* (GenBank accession number: IAAB01050803.1 ^19^) in the version 6 of the planarian “Dresden” transcriptome, publicly accessible on Planmine ^38^, identified as dd_Smed_v6_4796_0_1. *gfp* used as a negative control for the dsRNA was obtained as described in Pérez-Posada *et al*. ^15^, using the following primers: ggccgcggGACGTAAACGGCCACAATT (gfp-F), gccccggccGAACTCCAGCAGGACCATGT (gfp-R). All primers include linkers for Universal T7 primers. Primary PCR was performed using 1μL cDNA, 0.2μL Taq DNA polymerase (NEB), 2μL 10x buffer, 0.4μL dNTPs (2.5uM), 4μL forward primer (2.5μM), 4μL reverse primer (2.5μM) and 8.4μL H2O. The thermocycler program was 94°C (30 s); 35 cycles at 94°C (20 s), 55 °C (20 s) and 68°C (30 s); and 68°C (5 min).

The size and purity of PCR products was assessed in a 1% agarose gel. cDNA products of the expected sizes were cut under UV light and frozen in 20μL water. cDNA from the gel bands was used as template for the secondary PCR with Universal T7 primers. Secondary PCR was performed using 1μL of the primary PCR product, 0.2μL TAQ polymerase (NEB), 2μL 10x buffer, 0.4μL dNTPs (2.5μM), 0.5μL UniT7-F5 primer (10μM), 0.5μL UniT7-R5 primer (10μM) and 15.4μL water. The thermocycler program was 94°C (30 s); 5 cycles at 94°C (20 s), 50 °C (20 s) and 68°C (40 s); 35 cycles at 94°C (20 s), 65 °C (20 s) and 68°C (40 s); and 68°C (5 min).

**Cloning:** The secondary PCR product was cloned into a vector using NEB PCR cloning kit (NEB #E1202) following the manufacterer’s protocol. Briefly 1μL of the PCR product was ligated with 1μL of the linearized pMiniT 2.0 Vector for 12 min at room temperature followed by 2 min on ice. 1μL of the ligation mix was added to 50μL of competent cells which were transformed by heat shock at 42°C for 30 sec. *Escherichia coli* bacteria were plated on LB plates with ampicillin and incubated at 37°C overnight. Colonies were tested to check the size of the insert using an agarose gel.

**Miniprep:** Colonies with the right insert size were cultured in LB broth with ampicillin overnight at 37C and plasmid was retrieved by miniprep (NEB Monarch plasmid miniprep kit). The insert was retrieved by PCR using the following mix: 1μL TAQ polymerase (NEB), 10μL 10x buffer, 2μL dNTPs (2.5μM), 1μL UniT7-F5 primer (10μM), 1μL UniT7-R5 primer (10μM) and 84μL water. The thermocycler program was 94°C (30 s); 35 cycles at 94°C (20 s), 65°C (20 s) and 68°C (30 s); and 68°C (5 min). The PCR products were then purified with SPRI size selection (KAPA Pure Beads, Roche) following the manufacturer’s protocol. A small part of the purified PCR product was sent for sequencing to check the sequence of the insert. Products with the correct sequence were used as template for dsRNA synthesis.

**dsRNA synthesis:** dsRNA was synthesised by mixing 1 μg of purified cDNA, 12.5 μL of 2x Express Buffer (T7 RiboMAX, Promega), 2.5 μL of Express Mix (T7 RiboMAX, Promega), and up to 25 μL of nuclease-free water. The mix was incubated 4h at 37C. Remaining DNA was degraded by incubating with 2.5μL RNAse-free DNAse for 30 min at 37C. The reaction was stopped by adding 375μL of STOP solution (1M NH_4_OAc; 10mM EDTA; 0.2% SDS). The dsRNA was retrieved with phenol/chlorophorm extraction. 400μL of phenol/chlorophom pH 4.5 was added. After centrifugation, the supernatant was retrieved and 400μL chlorophorm was added. After centrifugation, the supernatant was retrieved and RNA was precipitated with 1mL cold ethanol. After 1 wash with 70% cold ethanol, the pelett was dried at 37°C and resuspended in 20μL water. The size of the product was checked with an agarose gel.

**dsRNA delivery:** For each RNAi condition (*cdh1*, *h2b*, gfp), 100 animals were collected and split equally in 2 replicates. dsRNA was delivered by feeding. 50μg of dsRNA were mixed in a food pellet made of 25μL of homogenized cow liver and 25μL of 1% low melting agarose. Animals were left to feed for 3h and the water was changed after feeding. 5 feedings with dsRNA were done over the course of 2 weeks. 4 days after the last feeding, phenotypes were scored. For each condition, 30 animals from each replicate were dissociated with ACME, 13 were cut to assess the effect of the knockdown on regeneration.

### SPLiT-Seq

SPLiT-Seq was performed as described in Emili *et al* ^39^, with a modified tagmentation protocol for the library L78 (*cdh1*/*h2b* RNAi). Library L47 (G1 and G2 enrichment) was tagmented with the Nextera DNA Library Preparation Kit from Illumina.

**Plate loading:** For L47, cells from 20 undissected untreated worms, dissociated in one ACME batch were distributed in 96 wells. For L78, all 3 samples (*cdh1*, *h2b* and *gfp* RNAi conditions, each with 2 replicates) of the experiment were loaded in the same RT plate (16 wells per sample) allowing processing without batch effect. They were loaded into specific wells of the 1^st^ round of barcoding, allowing separation of the reads during processing.

**FACS:** For L78, FACS sorting resulted in 7 sublibraries with 25k cells each. Due to a lower quality of the DNA after amplification, 1 sublibrary was not sequenced resulting in a dataset with 6 sublibraries. For L47, an additional gating step was added to split G1 and G2 cells into different tubes. L47.1 and L47.2 are enriched in G1 cells and have each 20k and 21k cells. L47.3 is enriched in G2 cells and has 15k cells. L47.4 has 20k cells which are a mix of G1 and G2 cells.

**PCR amplification:** All sublibraries were amplified for 8-9 qPCR cycles

**Tagmentation:** The sublibraries of L78 were tagmented using the Illumina DNA prep kit. 10 ng of DNA for each sublibrary were tagmented. PCR amplification was performed with the enhanced PCR mix from the kit. The primers used contain the i7 and i5 indexes used for sequencing. Each sublibrary receives a different i5 index. The following PCR program was used: 68°C (3 min); 98°C (3 min); 9 cycles at 98°C (45 s), 62 °C (30 s) and 68°C (2 min); and 68°C (1 min). Then 3 rounds of SPRI size selection were run: 0.5x keeping the supernatant to eliminate fragments bigger than 1000bp, 0.7x and 0.6x keeping the beads to eliminate fragments smaller than 500 bp.

### SPLiT-Seq data analysis

Sequencing, quality control, mapping and matrix generation was done as described in Emili e*t al*, 2024 ^39^. Mapping was done to the latest version of *Schmidtea mediterranea* genome ^33^. We used the genome annotation described in Emili *et al*, 2025 ^34^. The genes x cells matrices from the two experiments were loaded together in a Python Jupyter Notebook and we performed most of the analysis using scanpy ^40^. We filtered low quality cells (number of genes < 150).

**Doublet analysis:** We removed potential doublets using scrublet ^24^. We used an expected doublet rate of 0.1 and different thresholds for the two SPLiT-Seq libraries. For the G1G2 library, we used a threshold of 0.31 to identify doublets, and a threshold of 0.42 for the RNAi library. With those thresholds, 2.33% of the cells (4589 cells) were annotated as doublets and excluded from further analysis.

**Cell clustering, cell types identification and marker analysis:** We normalised the data using scanpy and selected 20 000 highly variable genes. We stored an adata.raw object at this stage that was later used for subclustering analyses and differential gene expression analysis. We then scaled the data using sc.pp.scale. Before building the knn graph, we integrated the two datasets with harmony. Then we built the kNN graph with the function sc.pp.neighbors with 30 neighbors and 110 pcs. A UMAP visualisation was generated (min_dist= 0.75, spread = 1.25) and the clustering was done using the Leiden algorithm, with resolutions ranging from 1 to 3 in steps of 0.5. We selected resolution 2.5 for annotation. Markers were calculated using the Wilcoxon and Logistic Regression methods. Cell type labels were assigned to cell clusters by visualising the expression patterns of known marker genes and integrating the data with published datasets using ingest with the dataset of Emili et al, 2024 ^39^ as reference. Broad cell types were obtained based on the dendrogram of cluster relatedness calculated by hierarchical clustering using sc.tl.dendrogram.

### Neoblast score

We selected the 50 top markers of the 2 main neoblasts clusters (neoblasts1 and neoblasts2) defined with the Wilcoxon method, making a total of 58 genes and calculated a ‘neoblast score’ for all cells. We then calculated the mean neoblast score per cluster and used it to annotate neoblasts and progenitors clusters. The early epidermal progenitors and neuronal progenitors are well described progenitors clusters. Any cluster with a mean neoblast score higher than the early epidermal progenitor clusters was considered as neoblasts (neoblast score > 0.258), any cluster with a mean neoblast score between those two was considered as progenitor or mixed progenitor cluster depending on the markers (neoblast score < 0.258, > 0.171), and any cluster with a neoblast score lower than the neuronal progenitors was considered as differentiated cluster (neoblast score < 0.171).

### MetaCell analysis

We performed an additional clustering using MetaCell ^26^. We ran the metacell using a custom python script as a wrapper. We set the target UMI number to 3000 UMIs per metacells, and defined the list of lateral genes as any gene linked to ribosomal biology on the genome annotation used in Emili et al and Pérez-Posada et al. ^15^. The resulting table was added to the adata object. Each metacell was labelled with the most frequent cell type label. We calculated the mean neoblast score of each metacell and assigned it to the high, medium, low neoblast score category (thresholds: 0.258 and 0.171).

### Cell abundances analysis

We used scCoda ^28^, a Bayesian model for compositional single-cell count data, to compare variations of cell abundances between clusters across the different treatments. Differential cell-type abundance between G1 and G2 samples was modelled using a Bayesian multinomial regression with condition as the explanatory variable (C(Cond, Treatment(’G1’))), using G1 as the reference condition. The same approach was used to compare GFP, CDH1, and H2B samples, with GFP as the reference condition (C(Cond, Treatment(’GFP’))). Posterior parameter estimation was performed using the default Hamiltonian Monte Carlo (HMC) sampling procedure implemented in scCODA. Credible changes in cell-type abundance were identified using the posterior inclusion probabilities after controlling the estimated false discovery rate (FDR) at 5% (est_fdr = 0.05). For visualization only, Fisher’s exact tests were additionally performed for each cell type to calculate log₂ odds ratios between conditions, which were used to illustrate the direction and magnitude of compositional changes but were not used for statistical inference.

### Subclustering analyses

We performed subsclustering analyses by slicing the adata object. For each subclustering analyses we started from the adata.raw object stored previously. For the subclustering of 4C cells we selected cells from the 4C sample only. For the subclustering of 4C cells with high neoblast score (NSc) we selected those cells of the 4C sample with NSc > 0.185. For the subclustering of 4C cells from neoblast cluster cells we selected those cells of the 4C from the broad group “neoblasts”. Then, for each condition, we subsampled randomly 1861 cells to prevent bias due to different cell numbers. Scaling, PCA and kNN graphs were built using the same parameters as the main analysis. Each subclustering analysis was clustered using resolutions ranging from 1 to 5 in steps of 1. Then, the identities of each cell in the main analysis were counted for each subcluster at each resolution, reporting clusters with at least 10% of cells belonging to one of the main analysis clusters. We calculated the average silhouette score ^29^ for each clustering resolution with the silhouette_score function from scikit-learn.

### Differential Gene Expression analysis

We performed DGE analysis using DESeq2 ^41^ using the adata.raw object stored previously. We generated independent pseudobulk matrices for each experiment to aggregate gene expression of cells belonging to the same replicate and i) for the RNAi dataset the same cell type and RNAi treatment, ii) for the FACS sorted G1 and G2 cells the same sorting gate. For the DGE analysis between FACS sorted G1 and G2 cells, we subset cells from neoblasts and progenitors clusters and with a neoblast score higher than 0.185. We randomly generated 2 pseudoreplicates and compared all G1 cells against all G2 cells. For the DGE analysis of the RNAi dataset, we used the 2 biological replicates and compared either *cdh1* or *h2b* RNAi against the *gfp* control condition. Genes with a p-value adjusted bellow 0.05 were identified as differentially expressed.

### OrthoFinder

We used OrthoFinder ^32^ to identify *S. mediterranea* orthologues of well-characterised cell cycle genes. We ran OrthoFinder with standard parameters using a set of 19 species spanning the major metazoan lineages and including *S. mediterranea* as well as well-annotated model species (*Capsaspora owczarzaki, Amphimedon queenslandica, Nematostella vectensis, Hofstenia miamia, Owenia fusiformis, Crassostrea gigas, Ramazzottius varieornatus, Adineta vaga, Bohtrioplana semperi, Dugesia japonica, Schmidtea polychroa, Schmidtea mediterranea, Drosophila melanogaster, Strongylocentrotus purpuratus, Branchiostoma lanceolatum, Lepisosteus oculatus, Danio rerio, Mus musculus, Homo sapiens*). In parallel, we gathered three different sources to identify cell cycle genes: a reference list of cell cycle-associated genes was compiled from *Homo sapiens*, *Mus musculus*, and *Drosophila melanogaster* based on the dataset from Dabydeen et al. ^42^ further expanded using genes annotated with cell cycle-related Gene Ontology (GO) terms in https://www.uniprot.org/UniProt ^43^. Any *S. mediterranea* gene falling within an orthogroup that contains cell cycle-regulated genes from these species was putatively assigned as cell cycle-regulated. In addition, we also labelled any *S. mediterranea* gene annotated with cell cycle-related functional categories according to eggNOG annotation ^44^.

Using this approach, we identified 1175 putative cell cycle genes. To determine which of those genes are expressed in the neoblasts, we preformed differential gene expression analysis between neoblasts and all other cell types using DESeq2. Among the 1175 identified cell cycle genes, 448 were significantly enriched in neoblasts. To assess potential gene loss and expansion events in *Schmidtea*, we generated a custom list of cell cycle genes of interest and compared it with the set of 1175 identified cell cycle genes.

## Acknowledgements

Research at the SCBE lab at Oxford Brookes University and at the Living Systems Institute is supported by MRC grants (MR/S007849/1 and MR/W017539/1), a BBSRC Grant (BB/V014447/1) and a Leverhulme Trust grant (RPG-2019-332 and RPG-2023-330) to JS. We thank Yuki Sato for providing the planarian sequences of *cdh1* and *cdc20*. OB was supported by a BBSRC DTP studentship from Oxford Brookes University and the University of Oxford. TS was supported by an A2i internship of the University of Exeter. We thank the technical team at Novogene for their expert technical support throughout this project. Flow cytometry was performed at the Sir William Dunn School of Pathology Flow Cytometry Facility, University of Oxford with the assistance of Dr Robert Hedley and Vasiliki Tsioligka. We thank Raif Yuecel at the Cytomics facility of the University of Exeter for help with cytometric analysis. We also thank all of the members of the SCBE lab for scientific discussions.

## Data availability

The datasets supporting the conclusions of this article are available in: GEO (scRNA-seq reads as well as processed files such as h5ad objects): GSE346035.

Code, including markdowns, functions, and scripts, for all analyses: https://github.com/scbe-lab/CellCycle_Schmidtea

## Author contributions

JS and SP conceived the study and designed the experiments. SP and AN performed RNAi, generated cell dissociations. SP generated single cell libraries. VM provided technical assistance and maintained animals. TS generated additional RNAi experiments. SP and OB performed single-cell analyses. APP and SP performed comparative orthology analysis. SP, OB, APP and JS generated the figures and wrote the manuscript, with contributions from all other authors. All authors read and approved the final version of the manuscript.

## Declaration of generative AI and AI-assisted technologies in the manuscript preparation process

During the preparation of this work, the authors used ChatGPT for language polishing and for assistance in generating and refining analytical code. The authors reviewed and verified the output of the AI-assisted analyses and edited the generated text and code as needed. The authors take full responsibility for the content of the published article and for the accuracy and integrity of the analyses and results.

## Supplementary Figures

**Figure S1:**
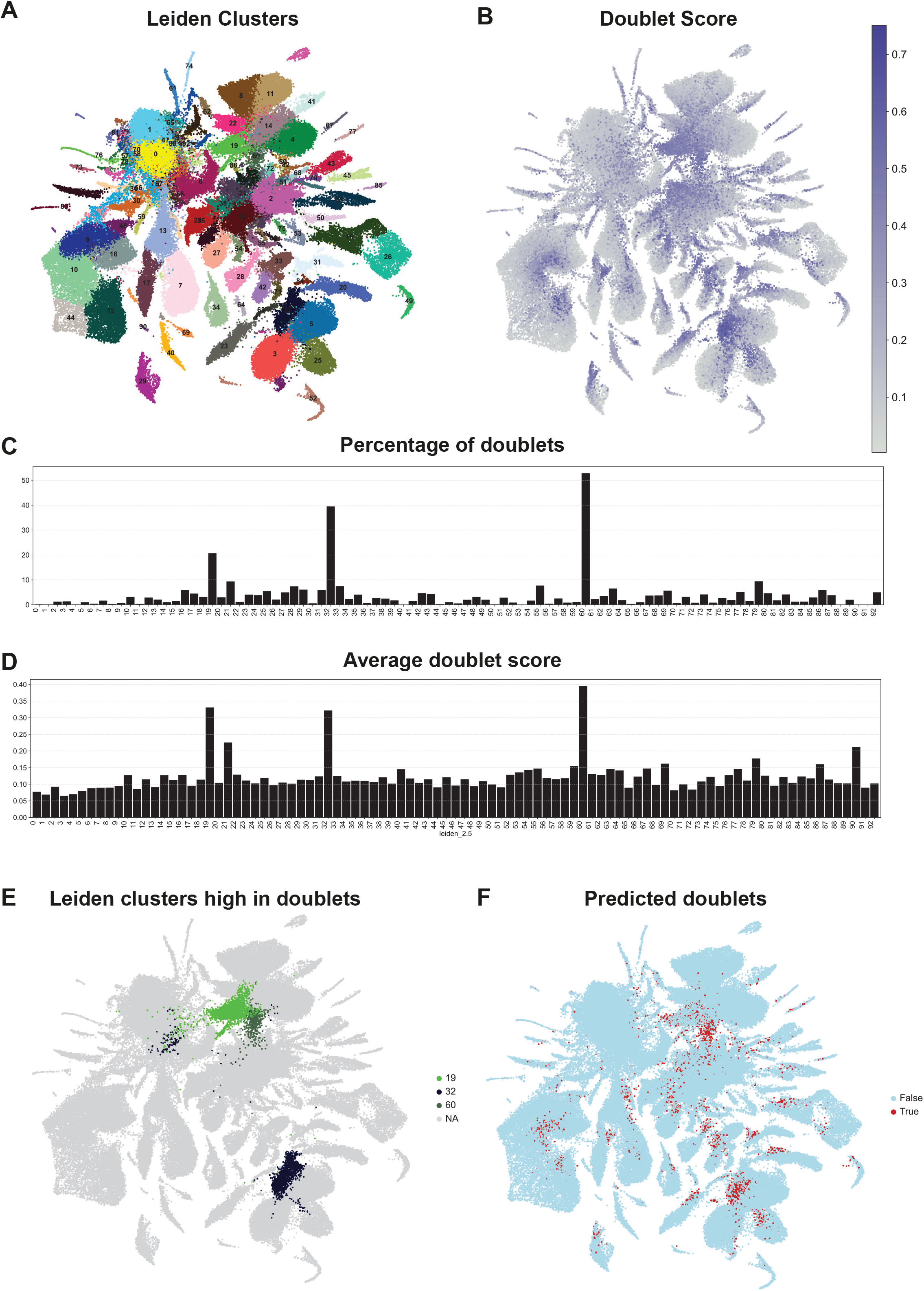
Doublet prediction using Scrublet. A. UMAP plot and clustering at the leiden 2.5 resolution of high-quality cells from the dataset (132037 cells with more than 150 genes) including doublets. B. UMAP plot showing the doublet score calculated using Scrublet. C. Percentage of cells marked as doublets per cluster. D. Average doublet score per cluster. E. UMAP plot highlighting the clusters containing more than 10% of doublets. F. UMAP plot highlighting the cells marked as doublets (doublet score > 0.42 for cells belonging to the FACS sorting experiment and > 0.31 for the cells belonging to the RNAi experiment).

**Figure S2:**
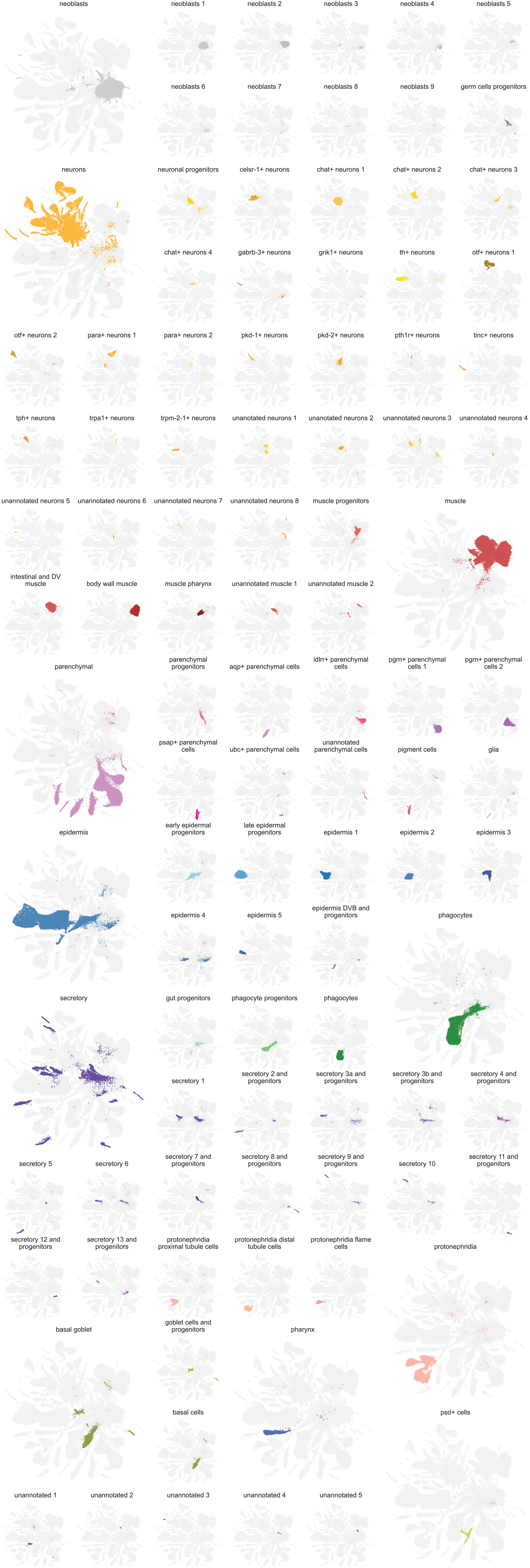
Visualisation of the annotated clusters. UMAP plot highlighting the 10 broad cell types and the 96 clusters (91 cell types and 5 unannotated) at the leiden 2.5 resolution.

**Figure S3:**
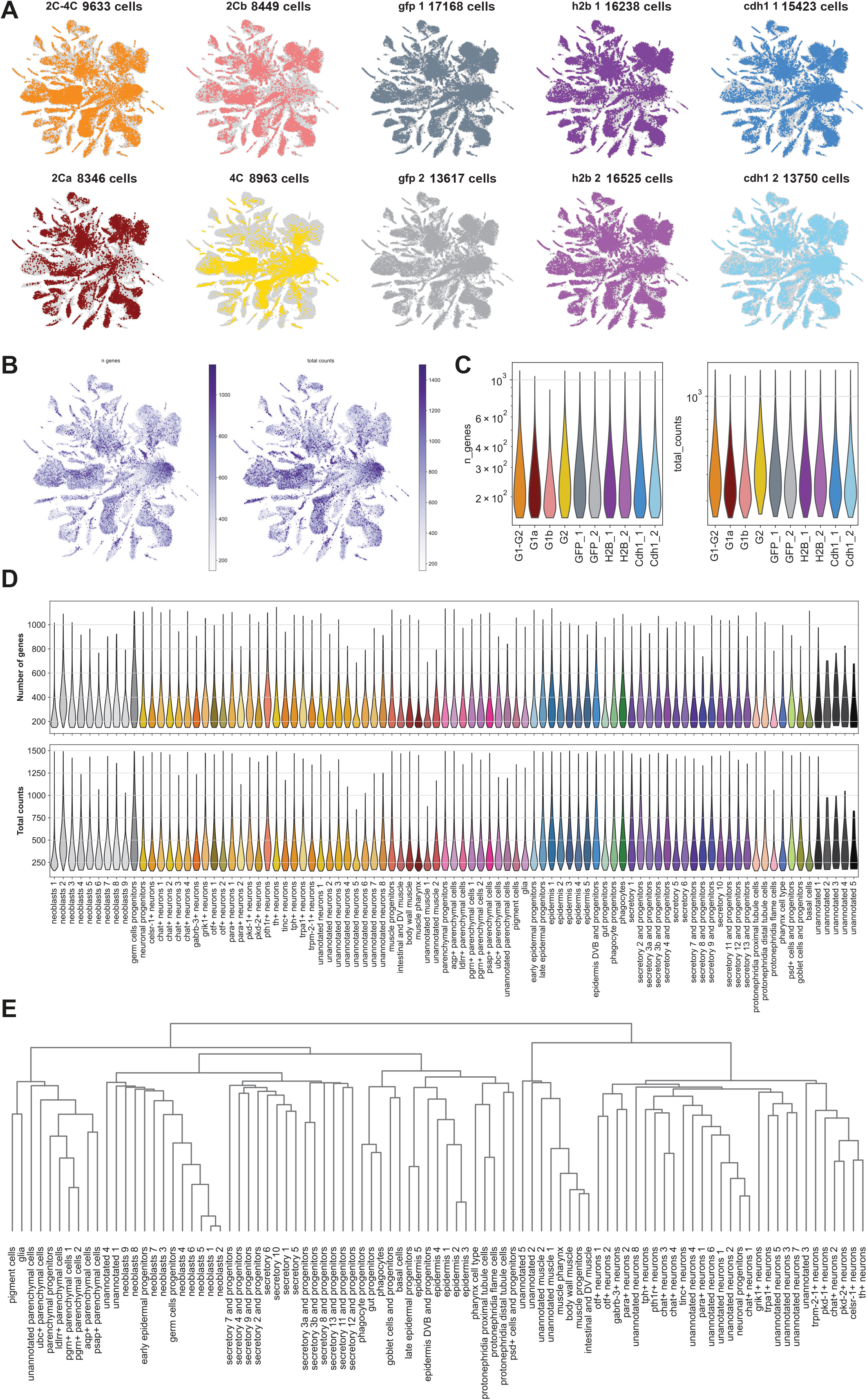
Sample distribution, quality metrics and cluster relatedness in the scRNA-seq dataset. A. UMAP plots highlighting the cells belonging to the 4 samples from the FACS sorting experiment (SPLiT-seq experiment 1: 2C-4C, 2Ca, 2Cb, 4C), and the 6 samples from the RNAi experiment (SPLiT-seq experiment 2: *gfp* 1, *gfp* 2, *h2b* 1, *h2b* 2, *cdh1* 1, *cdh1* 2). B. UMAP plot showing the gene numbers and UMI numbers per cell. C. Violin plots showing the distribution of the gene numbers and UMI numbers per cell for each sample. D. Violin plots showing the distribution of the gene numbers and UMI numbers per cell for each cluster. E. Dendrogram showing the relatedness of the clusters calculated by hierarchical clustering. This dendrogram was used to define the broad cell types.

**Figure S4:**
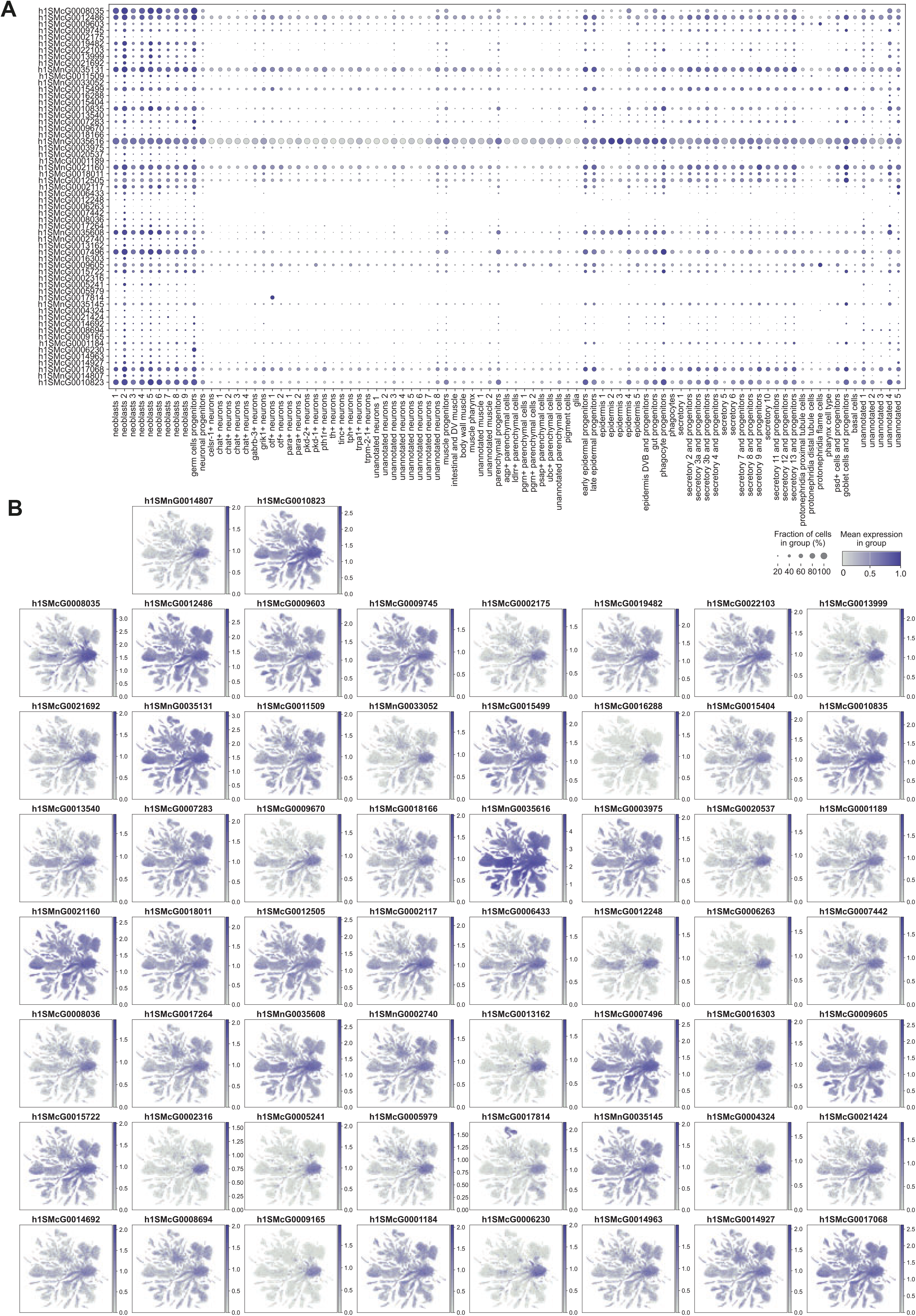
Expression patterns of the genes used to calculate the neoblast score. A. Dot plot showing the expression of the genes used for the neoblast score across all cell types. B. UMAP feature plots showing the expression patterns of the same genes.

**Figure S5:**
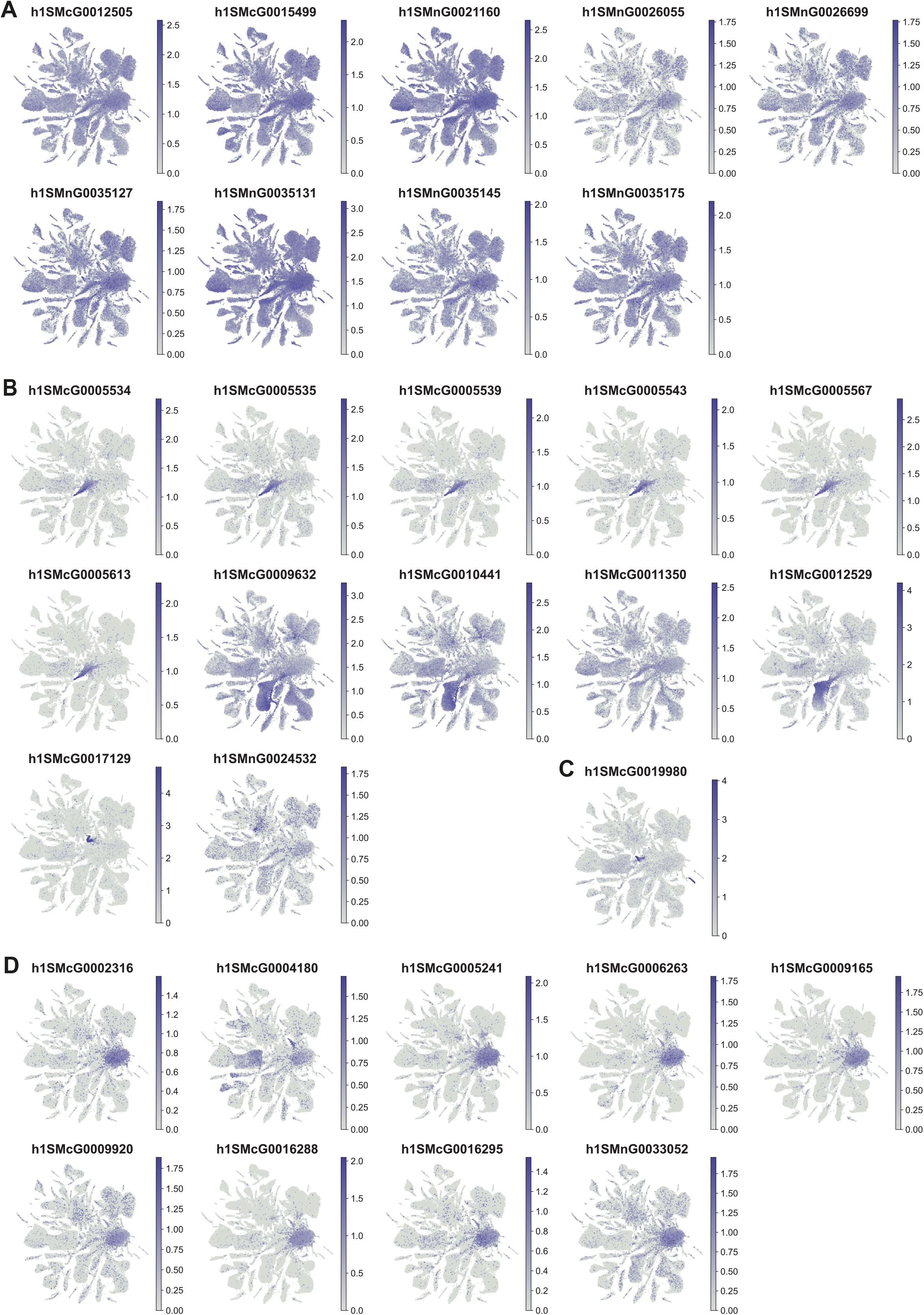
Expression patterns of differentially expressed genes between 2C and 4C neoblasts. A. UMAP feature plots showing the expression pattern of genes highly expressed in the neoblasts, upregulated in 2C neoblasts compared to 4C neoblasts. B. UMAP feature plots showing the expression pattern of genes with low expression in the neoblasts, upregulated in 2C neoblasts compared to 4C neoblasts. C. UMAP feature plot showing the expression pattern of a gene with low expression in the neoblasts, upregulated in 4C neoblasts compared to 2C neoblasts. D. UMAP feature plots showing the expression pattern of genes highly expressed in the neoblasts, upregulated in 4C neoblasts compared to 2C neoblasts.

**Figure S6:**
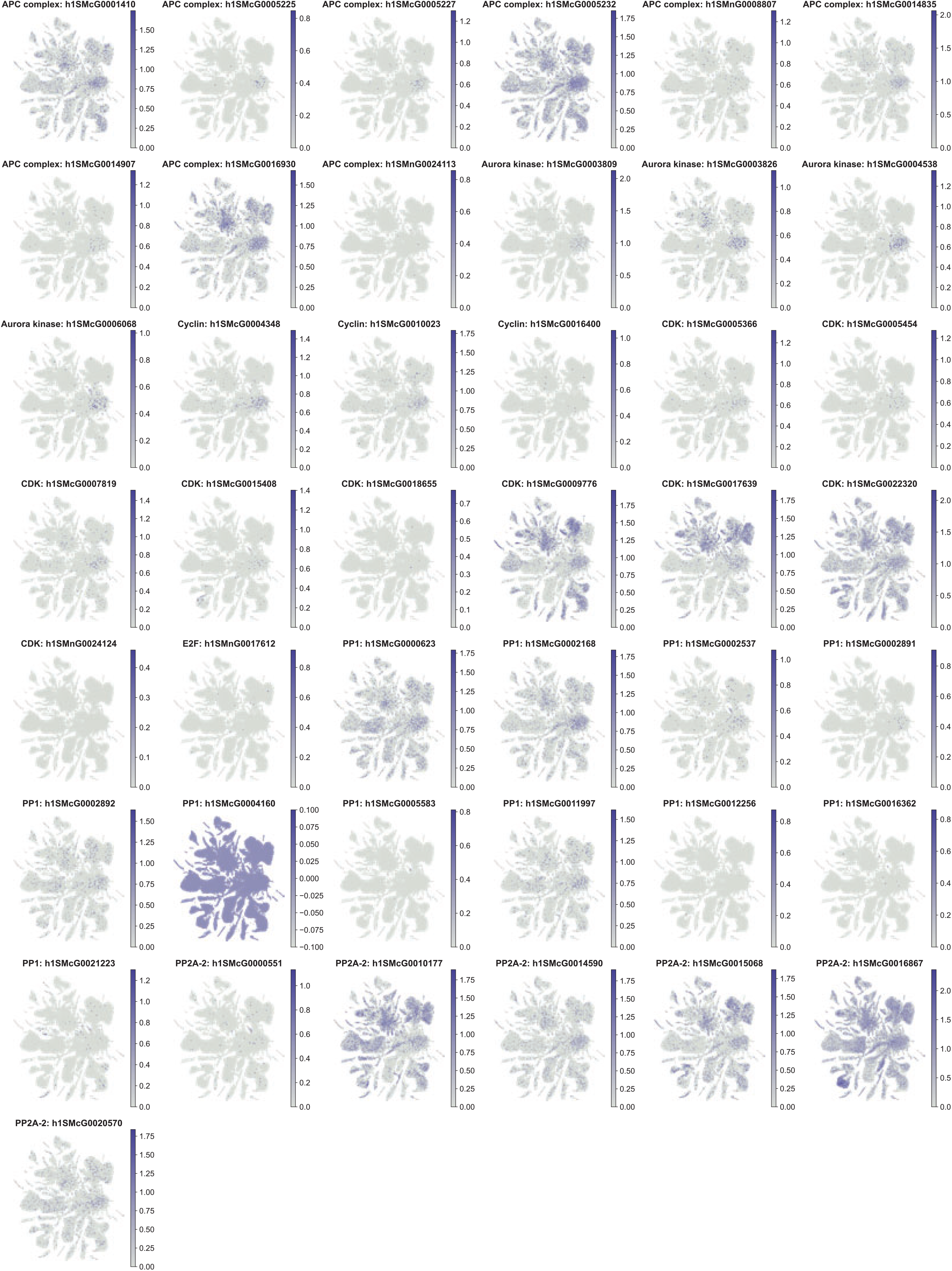
Expression patterns of cell cycle genes. UMAP feature plots showing the expression patterns of cell cycle genes identified using OrthoFinder.

**Figure S7:**
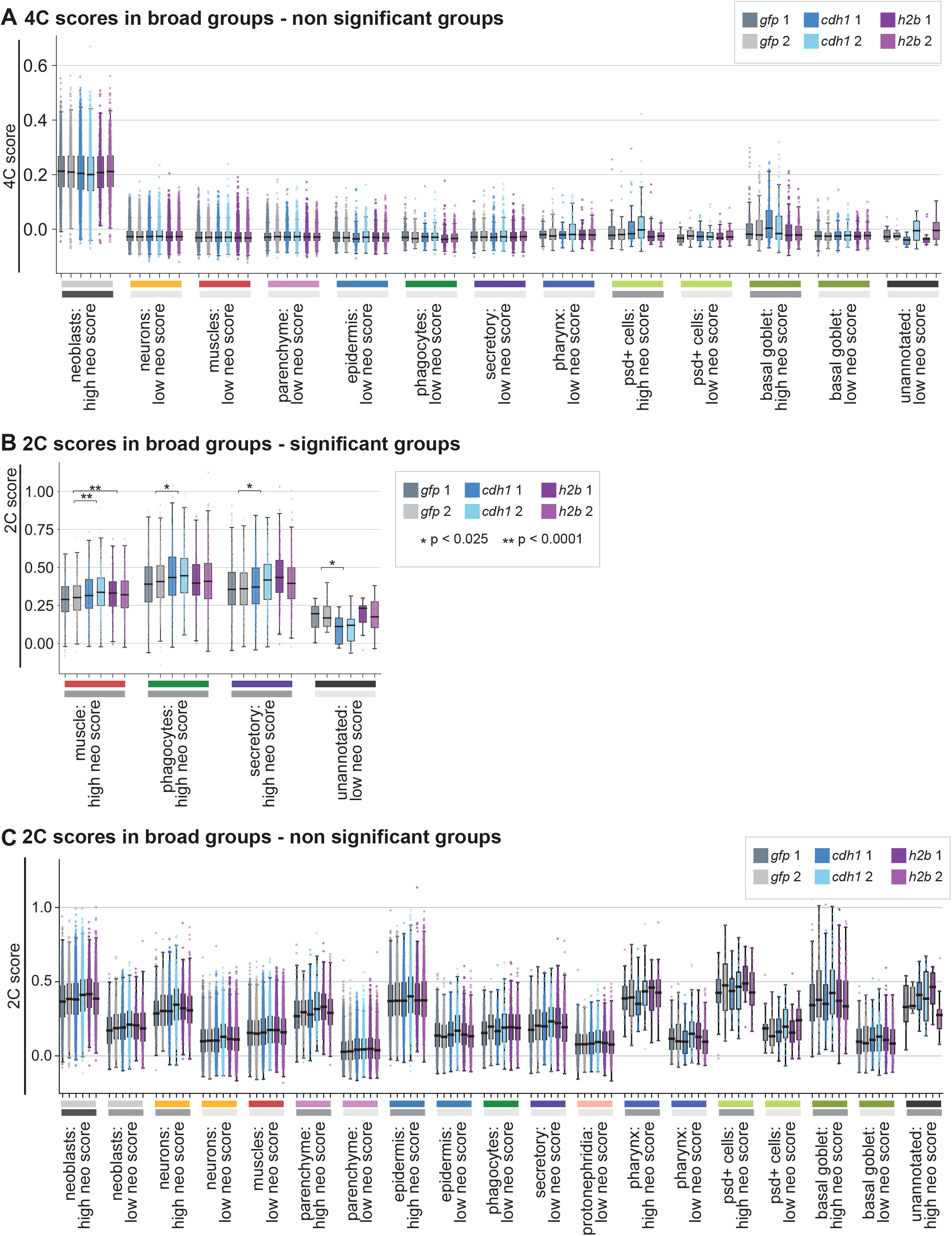
Differences in 4C and 2C scores across RNAi conditions in broad cell types. A–C. Box plots showing the 4C or 2C score for each RNAi treatment across broad cell type categories. Cell types were further classified into high- and low-neoblast-score groups based on their neoblast score. The asterisks indicate significant differences relative to the *gfp* RNAi control according to a linear mixed-effects model.

**Figure S8:**
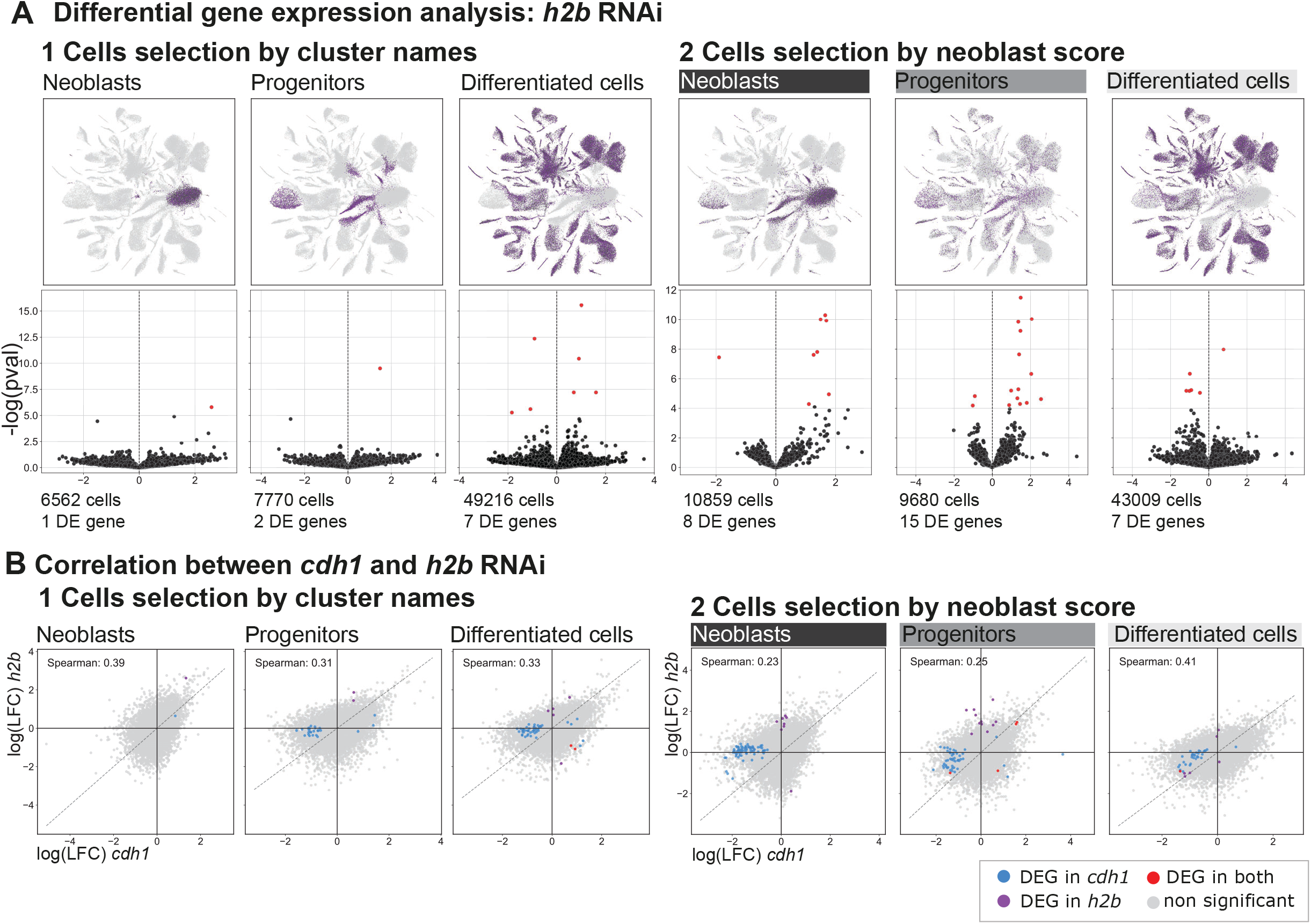
Distinct gene sets are affected by *cdh1* and *h2b* RNAi. A. Top: UMAP feature plots showing the cells selected for differential expression analysis. Bottom: Volcano plots highlighting in red genes significantly up- or downregulated in *h2b* RNAi relative to *gfp* RNAi. (1) Cells were grouped into neoblasts, progenitors, and differentiated cells according to cluster identity. (2) Cells were grouped into the same categories based on their neoblast score. B. Scatter plots comparing the differential expression changes induced by *cdh1* and *h2b* RNAi. Genes differentially expressed only in *cdh1* RNAi, only in *h2b* RNAi, or in both conditions are highlighted in blue, purple, and red, respectively.

## Data Files

**Data S1: Cluster Annotation**

Excel file with the cluster numbers for the leiden 2.5 resolution and the corresponding cell numbers in the different condition, cell type identity inferred from the ingest results and markers analysis and assigned colour.

**Data S2: Markers of the annotated cell types calculated by logistic regression.**

Excel file with the markers for each annotated cell type calculated with the Wilcoxon method. This file shows the gene id from the reference Schmidtea genome (Ivankovic *et al*), the equivalent id from the genome released by Grohme *et al*. and a name and description inferred from orthofinder.

**Data S3: Markers of the annotated cell types calculated by the Wilcoxon method.**

Excel file with the markers for each annotated cell type calculated with the Wilcoxon method. This file shows the gene id from the reference Schmidtea genome (Ivankovic *et al*), the equivalent id from the genome released by Grohme *et al*. and a name and description inferred from orthofinder.

**Data S4: Genes differentially expressed in 2C and 4C cells**

Excel file with the genes identified as differentially expressed in the 2C and 4C samples using DE

**Data S5: Cell cycle related genes identified using orthofinder**

List of all the cell cycle genes identified with orthofinder, including the ones not expressed in the neoblasts

**Data S6: *Schmidtea* has a simplified cell cycle regulation**

Excel file showing the results of a search for conserved cell cycle genes in the orthofinder results. This search highlights the fact that many genes are missing in *Schmidtea*

**Data S7: Cell cycle related gene families identified in *Schmidtea mediterranea* using orthofinder**

Excel file showing the gene families related to cell cycle regulation with orthologues in *Schmidtea mediterranea* identified using orthofinder

**Data S8: Cell cycle related genes used to calculate the cell cycle score**

List of the cell cycle genes identified with orthofinder expressed in the neoblasts. Those genes were used to calculate the cell cycle score

**Data S9: Genes differentially expressed in cells from the Cdh1 RNAi and H2B RNAi samples separating the cells according to the cluster names**

Excel file with the genes identified as differentially expressed in the *cdh1* and *h2b* RNAi samples using DESeq2. The cells were split into 3 categories (neoblasts, progenitors, differentiated cells) using the cell types annotation.

**Data S10: Genes differentially expressed in cells from the Cdh1 RNAi and H2B RNAi samples separating the cells according to the neoblast score**

Excel file with the genes identified as differentially expressed in the *cdh1* and *h2b* RNAi samples using DESeq2. The cells were split into 3 categories (neoblasts, progenitors, differentiated cells) using the neoblast score.

